# Natural phasic inhibition of dopamine neurons signals cognitive rigidity

**DOI:** 10.1101/2024.05.09.593320

**Authors:** Sasha C.V. Burwell, Haidun Yan, Shaun S.X. Lim, Brenda C. Shields, Michael R. Tadross

**Affiliations:** Department of Neurobiology, Duke University, Durham, NC; Department of Biomedical Engineering, Duke University, NC; Aligning Science Across Parkinson’s (ASAP) Collaborative Research Network, Chevy Chase, MD

## Abstract

When animals unexpectedly fail, their dopamine neurons undergo phasic inhibition that canonically drives extinction learning—a cognitive-flexibility mechanism for discarding outdated strategies. However, the existing evidence equates natural and artificial phasic inhibition, despite their spatiotemporal differences. Addressing this gap, we targeted a GABA_A_-receptor antagonist precisely to dopamine neurons, yielding three unexpected findings. First, this intervention blocked natural phasic inhibition selectively, leaving tonic activity unaffected. Second, blocking natural phasic inhibition accelerated extinction learning—opposite to canonical mechanisms. Third, our approach selectively benefitted perseverative mice, restoring rapid extinction without affecting new reward learning. Our findings reveal that extinction learning is rapid by default and slowed by natural phasic inhibition—challenging foundational learning theories, while delineating a synaptic mechanism and therapeutic target for cognitive rigidity.

## Main Text

To thrive, animals must predict and secure essential rewards, such as food and water. When predictions fail, persistence is crucial to overcoming temporary obstacles. However, excess persistence, known as perseveration, is generally maladaptive (*1–3*).

These processes are often studied using Pavlovian assays in which neutral cues are paired with appetitive rewards. Seminal work using this paradigm established that ventral tegmental area dopamine (VTA_DA_) neurons encode reward prediction error (RPE)—the difference between predicted and actual rewards (*4–9*). Investigating RPE’s synaptic origins (*10–12*) revealed that phasic excitation dominates when rewards exceed predictions, producing a burst in VTA_DA_ activity (positive RPE). As animals learn to predict rewards, phasic inhibition counteracts reward-evoked excitation (approaching zero RPE). Finally, when a prediction of reward fails, phasic inhibition becomes unopposed, yielding a pause in VTA_DA_ activity (negative RPE).

Regarding behavioral roles, the RPE framework has inspired many causal experiments delineating VTA_DA_ bursts and pauses as opposing forces. Artificial phasic excitation of VTA_DA_ neurons induces bursts that promote Pavlovian conditioning, reinforcing new cue-reward associations (*13–21*). Conversely, artificial phasic inhibition induces VTA_DA_ pauses that drive Pavlovian extinction, suppressing preexisting cue-reward associations (*22–24*).

However, widefield optogenetic perturbations tend to produce synchronous events that closely resemble natural bursts, where nearly all VTA_DA_ cells are recruited simultaneously by an unexpected external event (*25–27*). By contrast, natural VTA_DA_ pauses, signaling the failure of an internally generated prediction, exhibit irregular patterns across cells and time (*28*). In principle, these spatiotemporally distinct patterns could serve an essential role. For instance, synchronous events might broadcast globally to promote the formation of new synaptic engrams (*29*), whereas irregular patterns might act locally to update preexisting ones.

Nevertheless, the complexity of natural pauses also complicates their elimination. Widefield optical excitation cannot counteract natural phasic inhibition without producing bursts in cells that would have paused weakly or not at all (*28*). This concern is compounded by tonic-firing variability, ranging from 0.5 to 10 Hz (*5–7*), and the sensitivity of extinction learning to the intervention strength, being slowed by 20-Hz (*15, 30, 31*) but unaffected by 5-Hz (*31*) optogenetic excitation. While single-cell optogenetics presents a potential solution (*32, 33*), such precision has yet to be applied to VTA_DA_ cells.

In this study, we address the complexity of natural phasic inhibition by shifting the focus from VTA_DA_ pauses to their receptor-mediated origin. Specifically, we test the hypothesis that precisely blocking GABA_A_ receptors on VTA_DA_ cells would intercept natural phasic inhibitory inputs, slowing the rate of behavioral extinction. While broadly believed to be true, experimental support for this hypothesis has been indirect. Traditional GABA_A_ pharmacology (*34*) and optogenetic manipulations of GABA release in the VTA (*10, 35, 36*) support the canonical model, but with the caveat of affecting non-dopaminergic cells that directly impact reward learning (*37–40*). Knockout mice lacking the GABA_A_ ꞵ_3_ subunit in DA neurons exhibit normal extinction, but chronic compensatory changes were seen even in non-dopaminergic cells (*41*). To circumvent these issues, we used DART (drug acutely restricted by tethering), a technology enabling cell-specific, receptor-specific manipulations within minutes (*42, 43*).

### Gabazine^DART^ is a cell-specific, GABA_A_ receptor-specific antagonist

Our approach capitalizes on DART’s ability to precisely block native GABA_A_ receptors on VTA_DA_ neurons. To achieve cellular specificity, we use an AAV (adeno-associated viral vector) to express HTP (HaloTag Protein) exclusively on VTA_DA_ cells of DAT::Cre mice. HTP expression is stable and does not alter the physiology of VTA_DA_ cells (*43*). At a later time of interest, we apply gabazine^DART^, a two-headed ligand whose HTL (HaloTag Ligand) is efficiently captured by HTP, positioning the gabazine moiety to antagonize native GABA_A_ receptors only on VTA_DA_ cells (**Fig. 1A**). We co-deliver gabazine^DART^ with a small amount of Alexa647^DART^ to serve as a fluorescent proxy of drug delivery (**Fig. 1A**). Control mice are treated identically, except for use of a double-dead HTP (^dd^HTP), which cannot bind the HTL (**Fig. 1B**).

**Fig. 1:**
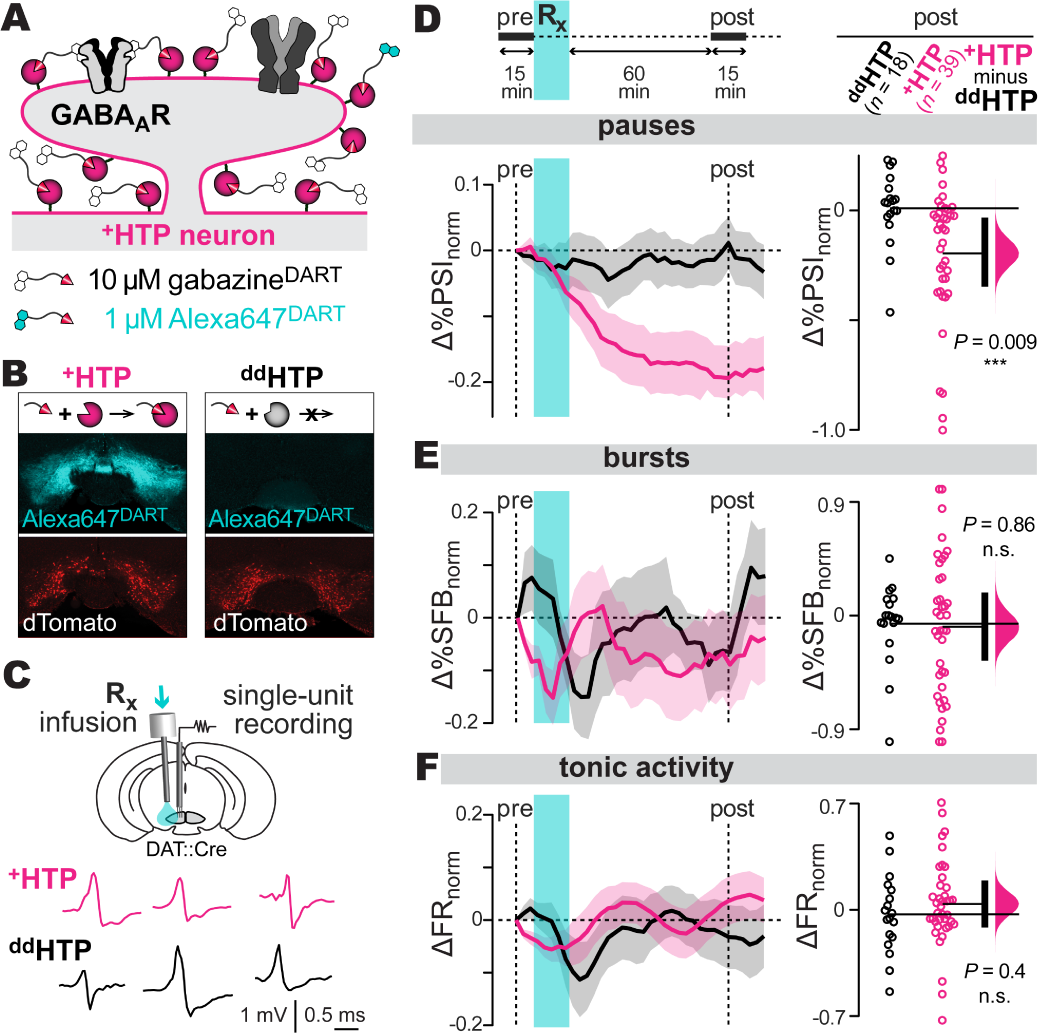
GABA_A_ receptors mediate VTA_DA_ pauses *in vivo* A**: DART technology.** AAV expression of the ^+^HTP protein (pink) enables cell-specific covalent capture of gabazine^DART^ (black) and Alexa647^DART^ (cyan) ligands. Once tethered, gabazine^DART^ blocks native GABA_A_ receptors, while Alexa647^DART^ enables fluorescent visualization of target engagement. B**: Example histology.** AAV expression of the active ^+^HTP or control ^dd^HTP in VTA_DA_ neurons is indicated by dTomato (red). All mice receive an intracranial ligand infusion of 10 µM gabazine^DART^ + 1 µM Alexa647^DART^, and are perfused 36 hr later for histology. Alexa647^DART^ (cyan) quantifies ligand target engagement. C**: Electrophysiology.** Top: an electrode bundle targeting the medial VTA enables *in vivo* extracellular recordings. A nearby cannula permits ligand infusion. Bottom: sample putative dopamine-neuron spikes, recorded in head-fixed animals, shown for ^+^HTP and ^dd^HTP mice. D**: Pauses in firing:** Top: time course of recording, baseline 15-min (pre-gabazine^DART^) followed by infusion and post-gabazine^DART^ recording. Bottom: pause metric, *%PSI* (percent of interspike intervals longer than twice the median interspike interval). Changes in *%PSI* compare a 15-min baseline (*%PSI*_pre_) to a 15-min sliding window (*%PSI*_post_) according to Δ*%PSI*_norm_=(*%PSI*_post_*-%PSI*_pre_)/(*%PSI*_post_+*%PSI*_pre_). Left: Δ*%PSI*_norm_ time course, mean ± SEM over cells (*n* = 18 ^dd^HTP cells, 3 mice; *n* = 39 ^+^HTP cells, 5 mice). Right: steady-state Δ*%PSI*_norm_ (1-hr post- gabazine^DART^) with individual cells (circles), group means (thin horizontal lines), mean-difference bootstrap (pink distribution), and 95% CI of the two-sided permutation test (vertical black bar); ^+^HTP and ^dd^HTP cells differ significantly (*P*=0.009). E-F**: Burst/tonic firing:** Analysis of *%SFB* (percent of spikes fired in bursts) and *FR* (firing rate) from the same cells; format as above.

In acute brain slices, dopamine neurons expressing ^dd^HTP were unaffected by gabazine^DART^, whereas neurons expressing the active ^+^HTP exhibited a rapid and nearly complete block of GABA_A_-mediated synaptic transmission (**Fig. S1A**) (*43*). Regarding receptor specificity, a saturating dose of tethered gabazine^DART^ did not alter excitatory glutamate receptors, nor did it impact the intrinsic pacemaker properties or action-potential waveforms of VTA_DA_ cells (**Fig. S1B-C**). Furthermore, Alexa647^DART^ did not influence VTA_DA_ physiology (**Fig. S1D**). Prior *in-vivo* tests found no behavioral effects of ambient gabazine^DART^ when infused into the VTA of awake ^dd^HTP mice (*43*). In ^+^HTP mice, gabazine^DART^ was tethered to VTA_DA_ neurons within minutes, with a single dose sufficing for two days of behavior (*43*). Together, these data validate tethered gabazine^DART^ as a precise, cell-specific antagonist of native GABA_A_ receptors.

### GABA_A_ receptors mediate natural phasic inhibition of VTA_DA_ neurons *in vivo*

In principle, GABA_A_ receptors could regulate various aspects of VTA_DA_ firing (*44*). We thus recorded VTA_DA_ action potentials in awake, head-fixed mice before and after delivery of gabazine^DART^ (**Fig. 1C**). We examined a panel of tonic, burst, and pause metrics using a sliding- window analysis spanning pre-gabazine^DART^ (15 min), drug infusion/equilibration (75 min), and post-gabazine^DART^ (15 min) periods. In ^dd^HTP mice, all metrics remained stable throughout the recording, indicating negligible effects of ambient gabazine^DART^ (**Fig. S2A**).

In comparing ^+^HTP mice to ^dd^HTP controls, we found that gabazine^DART^ substantially reduced the occurrence of spontaneous pauses. The effect is seen most readily in the main pause metric, *%PSI*, the percent of interspike intervals longer than twice the median interval. This metric was reduced by gabazine^DART^ in ^+^HTP mice relative to ^dd^HTP controls (two-sided permutation test, *P*=0.009, **Fig. 1D**). The effect was robust to adjustments in the definition of a pause, and could not be explained by a symmetrical change in interspike-interval variance (**Fig. S2B**). All other metrics exhibited no significant difference in ^+^HTP vs ^dd^HTP mice (**Fig. 1E-F**, **Fig. S2C-E**). Thus, the main effect of gabazine^DART^ is to prevent natural GABA_A_-mediated VTA_DA_ pauses from occurring. Histology confirmed ligand capture and specificity of viral expression: nearly all HTP expressing cells (99.7%) were dopaminergic and most dopaminergic cells in the VTA (∼64%) expressed HTP (**Fig. S2F-H**). We did not opto-tag cells given concerns that overexpression of a second membrane protein could hinder surface trafficking of HTP. Instead, we identified putative dopamine neurons, as others have (*45–47*), by their unique electrophysiological features (*48, 49*), criteria known to yield ∼12% false-positives (*49*).

To further scrutinize the data, we used our manipulation’s impact on pauses as a proxy for HTP expression and gabazine^DART^ capture on individual cells. Reductions in the number of pauses corresponded with a decrease in pause length (Pearson’s *r*^2^=0.28, *P*=0.0006, **Fig. S2I**), congruent with a pause-reduction effect. By contrast, no correlations were found with respect to burst parameters (**Fig. S2J**). Similarly, the median interspike interval showed no correlation to pause reduction (Pearson’s *r*^2^=0, *P*=0.95) while total spikes trended very weakly (Pearson’s *r*^2^=0.04, *P*=0.2, **Fig. S2K**). Thus, the underlying tonic rhythm was unaffected, while subtle total-spike increments are an expected proxy of pause reduction itself.

The absence of even transient changes in tonic firing suggests homeostatic processes operating faster than DART’s 15-min onset, consistent with dynamic setpoint regulation by A-type potassium channels (*50*). By contrast, millisecond phasic GABA signaling requires postsynaptic GABA_A_ receptors. Thus, gabazine^DART^ selectively blocks phasic inhibition of VTA_DA_ neurons while preserving tonic- and burst-firing characteristics.

### Blocking GABA_A_ receptors on VTA_DA_ cells accelerates Pavlovian extinction

Given that gabazine^DART^ prevents natural VTA_DA_ pauses, we expected it to slow extinction learning. We tested this in water-deprived mice, trained for 10 days to associate cue A (2.5 kHz tone, 1.5 sec) with sucrose-water reward in a head-fixed configuration (**Fig. 2A-B**). Anticipatory licking during the cue (prior to reward delivery) served as the primary learning metric (**Fig. 2C-D**; **Fig. S3A**). On day 11, mice received gabazine^DART^ and a 2 hr rest before resuming rewarded cue A trials for 15 min. Thereafter, Pavlovian extinction (unrewarded cue A trials) commenced, continuing into the final day (**Fig. 2E-F**, top). Contrary to expectations, we saw significantly accelerated extinction in ^+^HTP mice compared to ^dd^HTP controls (*P*=0.0043, two-sided permutation test, **Fig. 2G**)—opposite the hypothesized direction of influence.

**Fig. 2:**
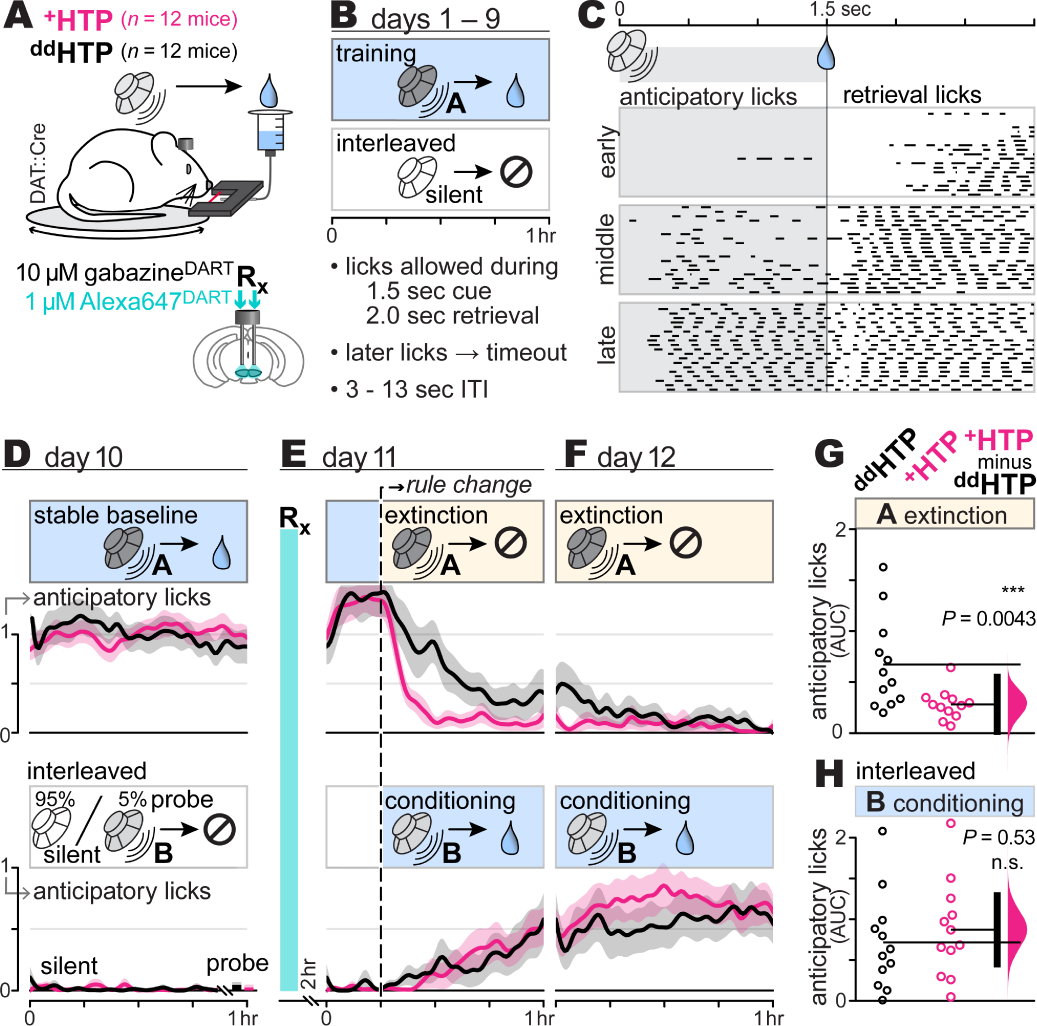
Blocking GABA_A_ receptors on VTA_DA_ cells accelerates Pavlovian extinction A**: Pavlovian behavior paradigm**. DAT::Cre mice with bilateral VTA cannula and ^+^HTP or ^dd^HTP expression in VTA_DA_ cells. Mice are head-fixed and presented with auditory cues and sucrose-water rewards. Licks detected with infrared beam; locomotion monitored via circular treadmill. B**: Training regimen**. Over 9 days, mice undergo sessions where cue A (2.5 kHz tone, 1.5 sec) reliably signals reward, interspersed with silent trials (no cue, no reward) to monitor nonspecific licking. Licking during the random (3-13 sec) inter-trial interval is discouraged with a timeout, training mice to ignore environmental sounds other than cue A. Each session lasts 1 hour and comprises 100-150 cue A trials. C**: Anticipatory licking**. Black line segments show beam breaks (licking) from a sample mouse on days 1 (early), 2 (middle), and 9 (late) of training. D**: Stable baseline**. To account for individual-mouse differences, day 10 anticipatory licking to cue A is calculated for each mouse and used as a constant of normalization for that animal. The same normalization constant is applied to cue A trials (top) and silent / cue B trials (bottom). Lines and shading are the normalized anticipatory licking mean ± SEM over mice (*n* = 12 ^dd^HTP mice; *n* = 12 ^+^HTP mice). E-F**: Pavlovian learning**. On day 11, mice receive gabazine^DART^ and a 2 hr rest. The first ∼15 min of the assay continue the prior day’s rules. Following the rule change, unrewarded cue A trials (extinction) are randomly interleaved with rewarded cue B trials (conditioning), contingencies which continue into day 12. Lines and shading are *lick*_norm_ mean ± SEM over mice (*n* = 12 ^dd^HTP mice; *n* = 12 ^+^HTP mice). G**: Extinction *AUC*** (area under the curve; licks_norm_ × hr), integrating *lick*_norm_ over 1.75 hr (post rule-change). *AUC* = 1.75 indicates no extinction (cue A anticipatory licking equal to that on day 10), while smaller values indicate greater extinction. *AUC* of individual mice (circles), group means (thin horizontal lines), mean- difference bootstrap (pink distribution), and 95% CI of the two-sided permutation test (vertical black bar) indicate a significant difference between ^+^HTP and ^dd^HTP mice (*P*=0.0043). H**: Conditioning *AUC***. Format as above, showing cue B conditioning trials from the same mice (*P*=0.53).

Mice were assayed during their dark (active) circadian phase, each being initially naïve to the task. To avoid frustration from complete reward denial, extinction trials of cue A were randomly interleaved with trials pairing a distinct cue B (11 kHz tone, 1.5 sec) with sucrose-water reward (**Fig. 2E-F**, bottom), thereby maintaining overall reward availability (*51*). This design also provided a within-mouse measure of Pavlovian conditioning, canonically driven by VTA_DA_ bursts, which we hypothesized would not be impacted by gabazine^DART^. Confirming this hypothesis, Pavlovian conditioning was unaffected by gabazine^DART^ (*P*=0.53, ^+^HTP vs ^dd^HTP, two-sided permutation test, **Fig. 2H**).

During training (days 1-10), we encouraged mice to ignore environmental sounds other than cue A by imposing a timeout for licking during the random (3-13 sec) inter-trial interval (**Fig. 2B**). This allowed cue B to remain novel (*52*), while achieving cue discrimination in the majority of mice: 89% (24 of 27) were unresponsive to rare probes of cue B presented on day 10, despite robust anticipatory licking to cue A, thereby satisfying our behavioral inclusion criteria (**Fig. S3B**).

Finally, after the session on day 12, we performed brain histology on every mouse to quantify target engagement. Locomotor enhancements, known to occur with VTA_DA_ disinhibition (*43, 53–56*), showed no correlation with gabazine^DART^ target engagement, nor with either form of Pavlovian learning (**Fig. S3C-E**). By contrast, a significant correlation between Pavlovian extinction and gabazine^DART^ target engagement was seen (Pearson’s *r*^2^=0.36, *P*=0.04, **Fig. S3F**), underscoring its dose-dependency.

### VTA_DA_ neural activity dynamics during Pavlovian behavior

Given these surprising behavioral findings, we further scrutinized the impact of gabazine^DART^ on VTA_DA_ neural dynamics during the Pavlovian assay. Photometry recordings were obtained from medial VTA_DA_ neurons that co-expressed HTP and jGCaMP8f (*57*), a cytosolic protein optimized to detect rapid calcium decrements (**Fig. 3A**). Histology confirmed fiber placement, AAV co- expression, and ligand capture (**Fig. 3B, Fig. S4A**). Of 18 mice that met behavioral criteria, 12 mice (6 experimental, 6 control) met a minimum signal-fidelity criterion (**Fig. S4B**).

**Fig. 3:**
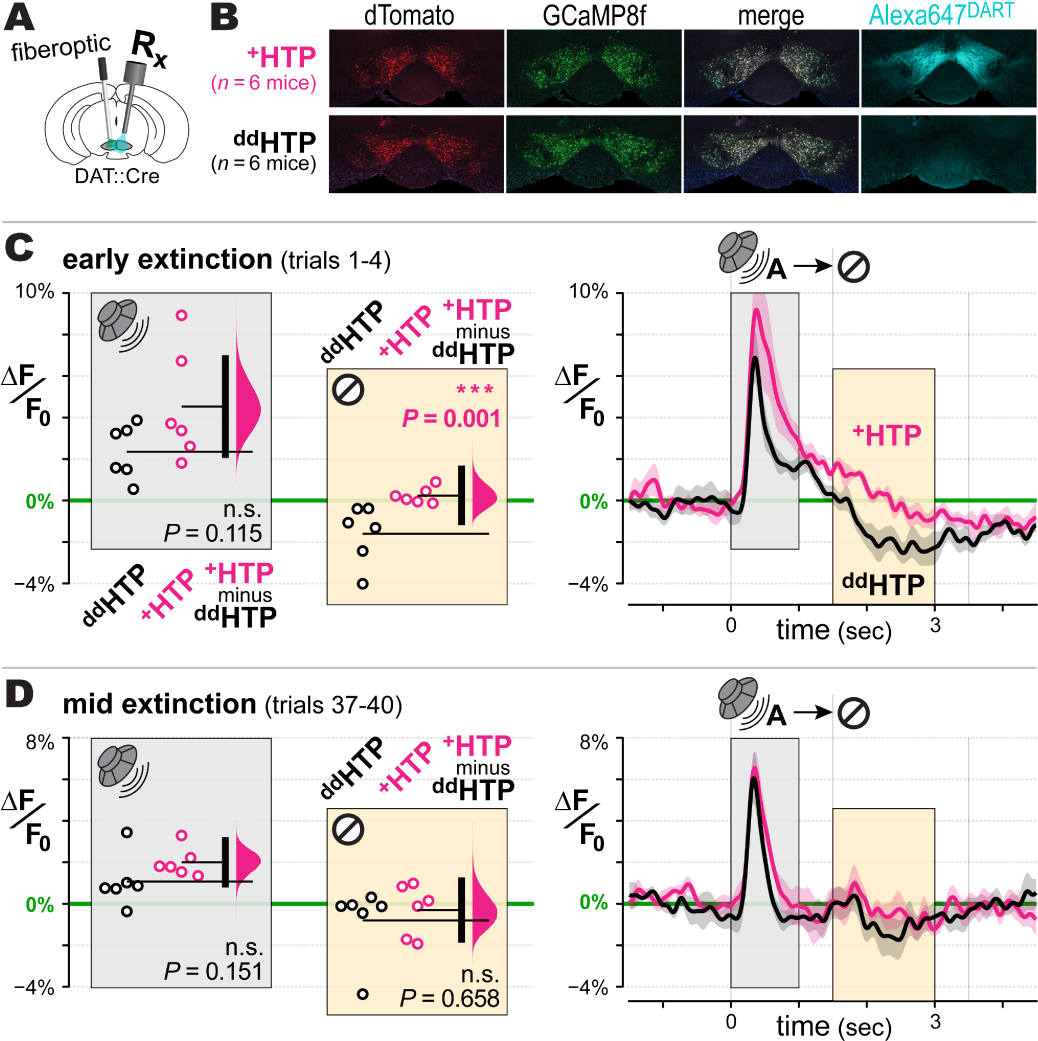
VTA_DA_ dynamics during Pavlovian behavior A**: Experimental setup**. DAT::cre mice injected with AAV-DIO-GCaMP8f and either AAV-DIO-^+^HTP_GPI_ or AAV- DIO-^dd^HTP_GPI_ in the VTA. Cannula and optical fiber implants permit intracranial DART infusions and calcium recording from VTA_DA_ neurons throughout the 12-day Pavlovian assay. B**: Example histology**. AAV expression of GCaMP8f (green) and the active ^+^HTP or control ^dd^HTP indicated by dTomato (red). All mice receive an intracranial ligand infusion of 10 µM gabazine^DART^ + 1 µM Alexa647^DART^, and are perfused 36 hr later for histology. Alexa647^DART^ (cyan) quantifies ligand target engagement. C**: Early extinction.** GCaMP8f responses in ^dd^HTP (black) vs ^+^HTP (pink) mice during the first four extinction trials. Right: time course of ΔF/F_0_ mean ± SEM over mice (*n* = 6 ^dd^HTP mice; *n* = 6 ^+^HTP mice). Analysis of ΔF/F_0_ during cue-burst (0 - 1 sec) and omission-pause (1.5 - 3 sec) is plotted in the left panel of corresponding color. Left: individual mice (circles), group means (thin horizontal lines), mean-difference bootstrap (pink distribution), and 95% CI of the two-sided permutation test (vertical black bar). ^+^HTP and ^dd^HTP mice were not statistically different during cue Burst (*P*=0.115), yet differed significantly during omission-pause (*P*=0.001). D**: Middle extinction**. Format as above, during extinction trials 37 – 40.

Control mice exhibited a canonical VTA_DA_ pause to reward omission (**Fig. 3C**, black data within yellow boxes). These pauses were prominent during early extinction trials and diminished quickly thereafter, aligning with prior studies (*31*). We thus focused on the first 4 extinction trials and observed the elimination of pauses in ^+^HTP/gabazine^DART^ mice (**Fig. 3C**, pink data within yellow boxes), with a significant difference from controls (two-sided permutation test, *P*=0.001, **Fig. 3C**). In subsequent trials, VTA_DA_ pauses diminished in control mice, becoming statistically indistinguishable from manipulated animals (*P*=0.658, **Fig. 3D**).

We saw no group differences before ligand infusion, nor post-gabazine^DART^ effects on baseline GCaMP signals (**Fig. S4C-E**), consistent with the stable tonic firing seen in our electrical recordings (**Fig. 1F**). Bursts to cue A were not significantly altered (**Fig. 3C-D**, **Fig. S4F-G**, gray boxes), also consistent with electrical recordings (**Fig. 1E**). As reported for unrewarded cues (*25*), a biphasic burst-pause to cue B appeared during probe (**Fig. S4E**) and early conditioning (**S4H,** top); this was unaffected by gabazine^DART^ suggesting no GABA_A_ involvement. During later conditioning, bursts to cue B trended larger but did not reach statistical significance (**Fig. S4H**, white boxes), consistent with unaltered behavioral conditioning (**Fig. 2H**).

Our data adhere to the canonical subtractive mechanism of RPE calculation (*10*), where phasic inhibitory predictions diminish bursts if the predicted reward is received (*P*=0.034, **Fig. S4F**, right inset) or produce a pause if the predicted reward is withheld (*P*=0.001, **Fig. 3C**). However, this correlative adherence to RPE belies a starkly different causal picture—one in which natural phasic inhibition of VTA_DA_ neurons favors persistence over adaptation (**Fig. 2**).

### Gabazine^DART^ impacts perseverative, but not flexible, mice

Mice exhibited variable rates of conditioning, which was anti-correlated with extinction in ^dd^HTP controls (Pearson’s *r*^2^=0.73, *P*=0.0004, **Fig. 4A**, black). We used conditioning, given its insensitivity to gabazine^DART^, to sort mice into upper and lower halves of this phenotypic spectrum. Slow-conditioning mice exhibited rapid cue A extinction (**Fig. 4B**, black), whereas fast- conditioning mice perseverated, responding to cue A despite repeated failure (**Fig. 4C**, black).

**Fig. 4:**
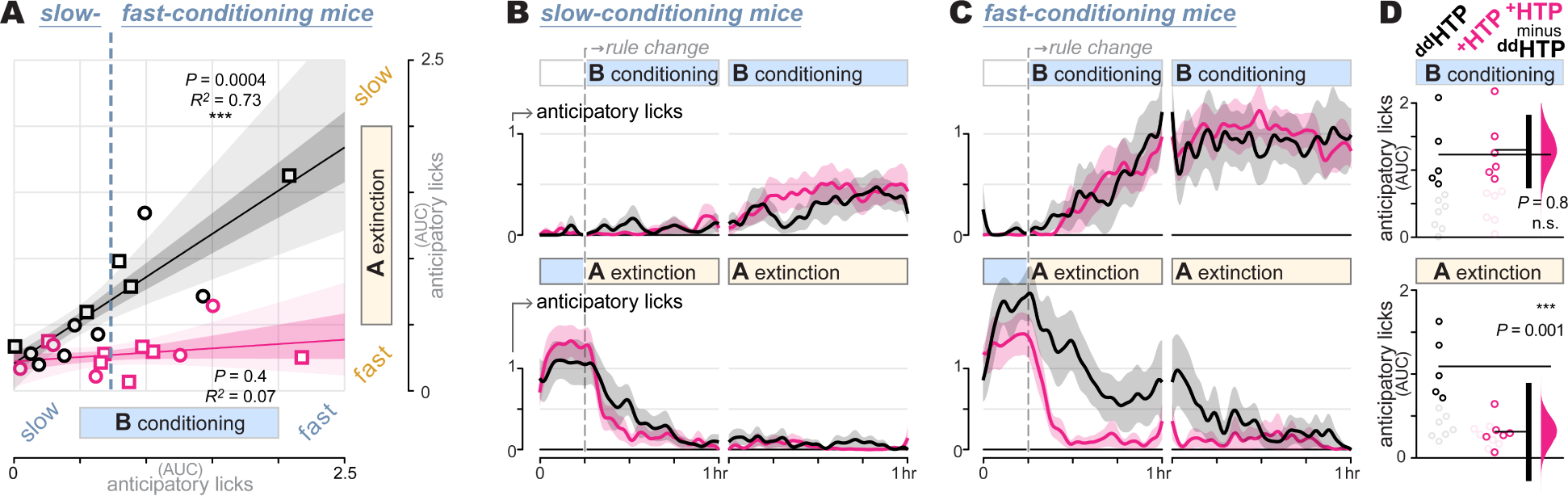
Gabazine^DART^ impacts perseverative, but not flexible, mice A**: Phenotypic spectrum.** Extinction-*AUC* vs conditioning-*AUC* measured within-mouse. Individual mice (squares = males, circles = females), regression fit (line), and regression 95% and 68% CI (light and dark shading) are shown for ^dd^HTP (black, *n* = 12) and ^+^HTP (pink, *n* = 12) mice. In ^dd^HTP mice, we observe an anti-correlation wherein fast-conditioning and slow-extinction (upper-right) tend to co-occur (Pearson’s *r*^2^ = 0.73, *P* = 0.0004). In ^+^HTP mice, the full spectrum of conditioning is seen, however extinction is uniformly rapid, independent of conditioning (Pearson’s *r*^2^ = 0.07, *P* = 0.4). Vertical dashed line illustrates the conditioning boundary (*AUC* = 0.75) chosen to divide mice into the approximate lower and upper halves of the phenotypic spectrum. B: **Slow-conditioning mice**. Analysis of mice exhibiting slow conditioning (*AUC* < 0.75), with conditioning (top) and extinction (bottom). Lines and shading are normalized anticipatory licking, mean ± SEM over mice (*n* = 7 ^dd^HTP mice; *n* = 6 ^+^HTP mice); C**: Fast-conditioning mice**. Analysis of mice exhibiting fast conditioning (*AUC* > 0.75), with conditioning (top) and extinction (bottom). Lines and shading are normalized anticipatory licking and mean ± SEM over mice (*n* = 5 ^dd^HTP mice; *n* = 6 ^+^HTP mice). D**: Fast-conditioning *AUC***. Summary data with the slow-conditioning subset of mice removed (faded circles), allowing a focused analysis of the fast-conditioning subset (*AUC* > 0.75). *AUC* of individual mice (circles), group means (thin horizontal lines), mean-difference bootstrap (pink distribution), and 95% CI of the two-sided permutation test (vertical black bar) indicate that ^+^HTP and ^dd^HTP differ significantly with regard to extinction learning (*P*=0.001).

Gabazine^DART^ eliminated perseveration in the fast-conditioning subset of mice, selectively accelerating extinction (*P*=0.001, two-sided permutation test, **Fig. 4D**, bottom) without altering their naturally fast conditioning (*P*=0.8, **Fig. 4D**, top). When administered to the other phenotypic category of mice, gabazine^DART^ had little impact on either form of Pavlovian learning (**Fig. 4B**). Overall, gabazine^DART^ caused extinction to become uniformly rapid across the phenotypic spectrum, eliminating its anti-correlation with conditioning (Pearson’s *r*^2^=0.07, *P*=0.4, **Fig. 4A**, pink). Consequently, a unique phenotype not seen in the control population emerged—mice adept at both rapid conditioning and rapid extinction (**Fig. 4C**, pink).

We did not observe trending sex differences (**Fig. 4A**), nor patterns in randomly interleaved trials that could explain phenotypic differences (**Fig. S5A**), consistent with previously reported intrinsic trait variability (*58, 59*). Initial training rates, which can be obscured by variability in task familiarization, did not predict later phenotypes (**Fig. S5B**). By contrast, we consistently observed the same phenotypic anti-correlation across a larger set of 25 mice pooled from the control arms of ongoing studies (**Fig. S5C**). This underscores the unique rapid-conditioning / rapid-extinction phenotype produced by VTA_DA_-specific GABA_A_ antagonism (**Fig. 4C**, pink).

## Discussion

The notion that learning is amplified by surprise is a central tenet in behavioral neuroscience. Canonically, surprise produces phasic dopamine signals that enhance cognitive flexibility, driving rapid learning, while tonic dopamine serves primarily as a baseline. Here, we reveal a fundamental inversion of these roles. Our finding that tonic dopamine allows rapid extinction learning aligns with a recent study suggesting that animals exhibit a default learning rate in the absence of phasic dopamine fluctuations (*21*). However, their focus on phasic excitation only allowed for transient increases above this default learning rate. Our study extends these findings by revealing that natural phasic inhibition can transiently slow the learning rate well below its default. Rather than conveying surprise, which enhances cognitive flexibility, we propose that natural phasic inhibition acts as a cognitive-rigidity signal, triggering skepticism toward new evidence that conflicts with prior expectations.

Two technical considerations arise from our findings. First, while our approach is specific to GABAergic over glutamatergic inputs, it remains unclear which subset of GABAergic inputs encodes the cognitive-rigidity signal. Given the importance of postsynaptic VTA_DA_ specificity to our findings, new tools must be developed to maintain this feature while refining input specificity—a constraint beyond the capability of existing tools (*43*). Second, since gabazine^DART^ and excitatory optogenetics both counteract phasic inhibition of VTA_DA_ neurons, their opposite behavioral effects merit discussion. Early studies using 20-Hz optogenetic excitation to overpower natural phasic inhibition induced artificial bursts, driving conditioning that could be mistaken for slowed extinction. Efforts to mitigate this confound by using 5-Hz excitation still induce artificial bursts that could obscure measures of extinction, consistent with the reported lack of behavioral effects (*31*). Thus, despite similar population-average effects, adding an artificial signal is not the same as blocking a natural one.

The sensitivity of behavior to VTA_DA_ activity patterns suggests that the brain could employ a vocabulary of patterns for different forms of learning. For instance, optogenetic inhibition of VTA_DA_ cells (*22–24*) resembles mildly aversive air puffs (*10–12*): both are unexpected external events that induce synchronous phasic inhibition of VTA_DA_ neurons, without the need for prior training. By contrast, our study examines phasic inhibition tied to appetitive reward omission, which exhibits complex spatial and temporal patterns (*28*), stems from distinct presynaptic origins (*11, 12, 60*), and requires prior training (*31, 61*). These two patterns of phasic inhibition may engage different dopamine-dependent plasticity rules (*31, 62–64*) with distinct objectives. Synchronous pauses may signal aversive surprise (*10–12, 65*), enhancing cognitive flexibility to predict and avoid future air puffs. Irregular pauses may signal cognitive rigidity—driving persistence despite failure.

A persistence mechanism can be adaptive in moderation (*31, 61*), but may become maladaptive, leading to perseveration as seen in schizophrenia (*66*), obsessive-compulsive disorder (*67*), addiction (*68*), and Parkinson’s disease (PD) (*69*). While many of these disorders have been characterized as hyper- or hypo-dopaminergic, our study shows that subtle changes in dopamine-neuron activity can have an outsized effect on behavior. This raises the question of whether modulating specific phasic components of dopamine signaling could offer a better therapeutic profile than current treatments that modulate dopamine signaling globally and continuously. While much remains to be done, the precision of gabazine^DART^, both in its specificity for extinction over conditioning behaviors and for perseverative over non-perseverative individuals, underscores the potential of targeted synaptic interventions for treating neurological disorders within a diverse population (*70*).

## Acknowledgements

We would like to thank Isaac Weaver and Janani Sundararajan for help designing and building the behavioral assay; Ankit Choudhury for performing tyrosine hydroxylase immunostaining; Konstantin Bakhurin for providing training on implanting electrodes into the VTA; and Robin Blazing for providing training on spike sorting. We also thank Rene Carter, Rich Mooney, Nicole Calakos, Mark Harnett, Elias Issa, Steve Lisberger, Erin Calipari, Vijay Namboodiri, and Josh Dudman for insightful feedback on the manuscript. This work was supported by Duke University Startup Funds (MRT), NIH grants RF1-MH117055 and DP2-MH1194025 (MRT), and by the joint efforts of The Michael J. Fox Foundation for Parkinson’s Research (MJFF) and the Aligning Science Across Parkinson’s (ASAP) initiative. MJFF administers grant ASAP-020607 on behalf of ASAP and itself.

## Author Contributions

See **Table S2** for detailed author contributions. **Conceptualization:** SCVB, MRT. **Methodology:** SCVB, HY, SSXL, BCS, MRT. **Software:** SCVB, MRT. **Validation:** SCVB, HY, SSXL, BCS, MRT. **Formal Analysis:** SCVB, HY, MRT. **Investigation:** SCVB, HY. **Resources:** HY, SSXL, BCS, MRT. **Data Curation:** SCVB. **Writing, Original Draft:** SCVB. **Writing, Review and Editing:** SCVB, HY, SSXL, BCS, MRT. **Visualization:** SCVB, HY, MRT. **Supervision:** MRT. **Project Administration:** SCVB, BCS, MRT. **Funding Acquisition:** MRT.

## Competing Interests

MRT and BCS are on patent applications describing DART. Other authors declare no competing interests.

## IP Rights Notice

For the purpose of open access, the author has applied a CC-BY public copyright license to the Author Accepted Manuscript (AAM) version arising from this submission.

## Data and Software Availability

All data and software are publicly available.

## Protocols

https://doi.org/10.17504/protocols.io.j8nlk8ekdl5r/v1

## Software

https://github.com/tadrosslab/VTA_GABA_paper and https://doi.org/10.5281/zenodo.10951255

## Datasets

- **Fig. 1, Fig. S1-2:** https://doi.org/10.5281/zenodo.10904059
- **Fig. 2,4, Fig. S3, 5:** https://doi.org/10.5281/zenodo.10903566 and https://doi.org/10.5281/zenodo.10908572
- **Fig. 3, Fig. S4:** https://doi.org/10.5281/zenodo.10908502

## Contact for Reagent and Resource Sharing

Further information and requests for resources and reagents should be directed to and will be fulfilled by the corresponding author Michael R. Tadross, MD, PhD.

## METHODS

### Mice

DAT-IRES-Cre (Jackson Labs 006660) mice were group housed by age and sex (max 5 per cage) in a standard temperature and humidity environment. For breeding, mice were housed under a normal 12-hr light/dark cycle and with food and water provided *ad libitum*. Experimental mice were transitioned to reverse-light-cycle and water-restriction conditions, as detailed below. All experiments involving animals were approved by the Duke Institutional Animal Care and Use Committee (IACUC), an AALAC accredited program registered with both the USDA Public Health Service and the NIH Office of Animal Welfare Assurance, and conform to all relevant regulatory standards (Tadross protocols A160-17-06, A113-20-05, A091-23-04).

### Recombinant Adeno-associated Viral (rAAV) Vectors

All custom viral vectors were produced by the Duke Viral Vector Core or VectorBuilder, kept frozen at -80°C until use, then diluted to the desired titers using sterile hyperosmotic PBS and kept at 4°C for up to 4 weeks.

### Acute Brain Slice Electrophysiology

DAT-IRES-Cre mice (5 females, 3 males, 8-10 weeks) were anesthetized and stereotaxically injected with 400 nL of either AAV_rh10_-CAG-DIO-^+^HTP_GPI_- 2A-dTomato-WPRE or AAV_rh10_-CAG-DIO-^dd^HTP_GPI_-2A-dTomato-WPRE (2 × 10^12^ VG/mL, 100 nL per site, two tracks with two depths per track: -3.2 mm AP, ±0.5 mm ML, -5.0/-4.5 mm DV) using a custom Narishige injector. After 3-5 weeks for expression, mice were deeply anesthetized with isoflurane and euthanized by decapitation. Coronal brain slices (300 µm) containing VTA were prepared by standard methods using a Vibratome (Leica, VT1200S), in ice-cold high sucrose cutting solution containing (in mM): 220 sucrose, 3 KCl, 1.25 NaH_2_PO4, 25 NaHCO_3_, 12 MgSO_4_, 10 glucose, and 0.2 CaCl_2_ bubbled with 95% O_2_ and 5% CO_2_. Slices were then placed into artificial cerebrospinal fluid (aCSF) containing (in mM): 120 NaCl, 3.3 KCl, 1.23 NaH_2_PO_4_, 1 MgSO_4_, 2 CaCl_2_, 25 NaHCO_3_, and 10 glucose at pH 7.3, previously saturated with 95% O_2_ and 5% CO_2_. Slices were incubated at 33°C for 40-60 min in bubbled aCSF and allowed to cool to room temperature (22-24°C) until recordings were initiated.

Recordings were performed on an Olympus BX51WI microscope, where slices were perfused with bubbled aCSF at 29-30°C with a 2 ml/min flow rate. To isolate GABA_A_ IPSCs, the external solution was supplemented with DNQX (20 µM, AMPA antagonist) and AP-V (50 µM, NMDA antagonist). Alternately, to isolate AMPA-mediated EPSCs, aCSF was supplemented with picrotoxin (50 µM, GABA_A_R antagonist) and AP-V (50 µM). Finally, NMDA-mediated EPSCs were isolated with picrotoxin (50 µM) and DNQX (20 µM).

For voltage-clamp, the internal solution contained (in mM): 135 CsCl, 2 MgCl_2_, 0.5 EGTA, 10 HEPES, 4 MgATP, 0.5 NaGTP, 10 Na_2_-phosphocreatine, and 4 QX314 (lidocaine N-ethyl bromide), pH 7.3 with CsOH (290 mOsm). For current-clamp, we used (in mM) 130 K-gluconate, 5 KCl, 2 MgCl_2_, 0.2 EGTA, 10 HEPES, 4 MgATP, 0.5 NaGTP, and 10 phosphocreatine, pH adjusted to 7.3 with KOH (290 mOsm). Internal solutions were used to fill glass recording pipettes (4-6 MΩ). The liquid junction potential, estimated to be 15.9 mV, was not corrected.

Whole-cell recordings were obtained with Multiclamp 700B and Digidata 1440A, which were controlled by pClamp 10.7 acquisition software (Molecular Devices). Signals were filtered at 10 kHz. A stimulating electrode was placed 60-100 μm from the recorded neuron. Evoked IPSC or EPSC signals were elicited by electrical stimuli of 0.3 ms duration and 150-300 μA (60-70% maximum responses), with a repetition interval of 15 sec. Our inclusion criteria required that cells maintain stable access and holding currents for at least 5 min. In particular, series resistance is monitored using 5–10 mV hyperpolarizing steps interleaved with our stimuli, and cells are discarded if series resistance changed more than ∼15% during the experiment. The stored data signals were processed using Clampfit 10.7 (Axon Instruments).

### *In Vivo* Electrophysiology Experiments

Adult DAT-IRES-Cre mice (2 females, 6 males; 12 -16 weeks old) were anesthetized and stereotaxically injected with 400 nL of either AAV_rh10_-CAG- DIO-^+^HTP_GPI_-2A-dTomato-WPRE, AAV_rh10_-CAG-DIO-^dd^HTP_GPI_-2A-dTomato-WPRE (2 × 10^12^ VG/mL), AAV_rh10_-CAG-CreON-W3SL-^+^HTP_GPI_-IRES-dTomato-Farnesylated, or AAV_rh10_- CAG-CreON-W3SL-^dd^HTP_GPI_-IRES-dTomato-Farnesylated (1 × 10^12^ VG/mL) (100nL per site, two tracks with two depths per track: -3.2 mm AP, ±0.5 mm ML, -5.0/-4.5 mm DV) with a custom Narishige injector. Mice were implanted with a single-drive movable micro-bundle electrode array (Innovative Neurophysiology, Inc.; 23 µm Tungsten Electrodes, 16 / bundle; 0.008” silver ground wire) above the left VTA (-3.2 mm AP, -0.5 mm ML, -4.0 mm DV). The silver ground wire was wrapped securely around two ground screws, one placed in the skull above the cerebellum and one above the right olfactory bulb. A unilateral metal cannula (P1Tech; C315GMN; cut to 13.5 mm) was implanted laterally adjacent to the electrode bundle (-3.2 mm AP, -1.3 mm ML, -4.0 mm DV). Mice were fitted with a plastic head bar adhered to the skull with OptiBond and dental cement. Mice were singly or pair housed post-surgery, in a 12-hr light/dark cycle, with food and water provided *ad libitum*. Pair-housed mice were outfitted with head hats that clip to specially designed head bars to prevent cannula or electrode damage from chewing by cage mates (*71*).

Electrophysiology recordings and DART infusions were performed at least 3 weeks after surgery to allow for recombinant protein expression. The electrode bundle was manually advanced three times: (1) 208 µm at least one week after surgery, (2) another 208 µm one week later, and (3) 104 µm one week later. This placed the electrodes at -4.5 mm DV, at the top of the VTA. After a few days for recovery, electrophysiological recordings were made with an Intan RHD 16-channel headstage with accelerometer (C3335) attached to an Open Ephys Acquisition Board via an Intan RHD 1-ft ultra-thin SPI interface cable (C3211). Data was collected using the Open Ephys GUI (*72*). Putative dopamine neurons were identified via their canonical features: tonic firing between 0 and 10 Hz, with bursting; wide biphasic or triphasic waveform; and large amplitude (*48, 49*). If no putative dopamine neurons were observed online, electrodes were advanced an additional 26-52 µm; this cycle was repeated until multiple channels with putative dopamine neurons were observed, at which point a recording was obtained.

DART ligands, stored as pure-compound aliquots, were freshly thawed on the day of use and dissolved in sterile artificial cerebrospinal fluid (aCSF) containing (in mM): 148 NaCl, 3 KCl, 1.4 CaCl_2_, 0.8 MgSO_4_, 0.8 Na_2_HPO_4_, 0.2 NaH_2_PO_4_. The final reagent solution contained 10 µM gabazine.7^DART.2^ + 1 µM Alexa647.1^DART.2^. This solution was loaded into an internal cannula designed to project 0.5 - 1.5 mm from the guide cannula, with progressively longer internals used on successive infusions. Mice were head-fixed, the internal cannula inserted, and the Innovative Neurophysiology electrode bundle was connected to the Intan headstage. After obtaining a 15 min baseline recording, we infused 1.5 µL of DART reagent over 15 min (0.1 µL/min; Harvard Apparatus PhD Ultra pump; 5 µL Hamilton syringe), and continued the recording (120 min total). After completion of the recording, electrodes were advanced 26-52 µm (*73*). Mice were given at least two weeks for recovery between recordings, which we have shown is sufficient to allow for complete HTP protein turnover (*43*).

Spike sorting of the raw data was performed using SpyKING CIRCUS, an open-access software package allowing for semi-manual spike sorting on multichannel extra-cellular recordings (*74*). Detection parameters included: spike threshold = 4; N_t (width of templates) = 2 or 3; peaks = positive. Filtering parameters used 250 Hz as the cutoff frequency for the Butterworth filter. All other parameters in the configuration file were standard as recommended by the SpyKING CIRCUS documentation. Only templates that matched all features of putative dopamine neurons and exhibited consistent spiking across the whole two-hour recording window were kept for analysis. All semi-manual spike sorting and template extraction were performed by SCVB for consistency.

Custom MATLAB code was used to extract the following metrics:

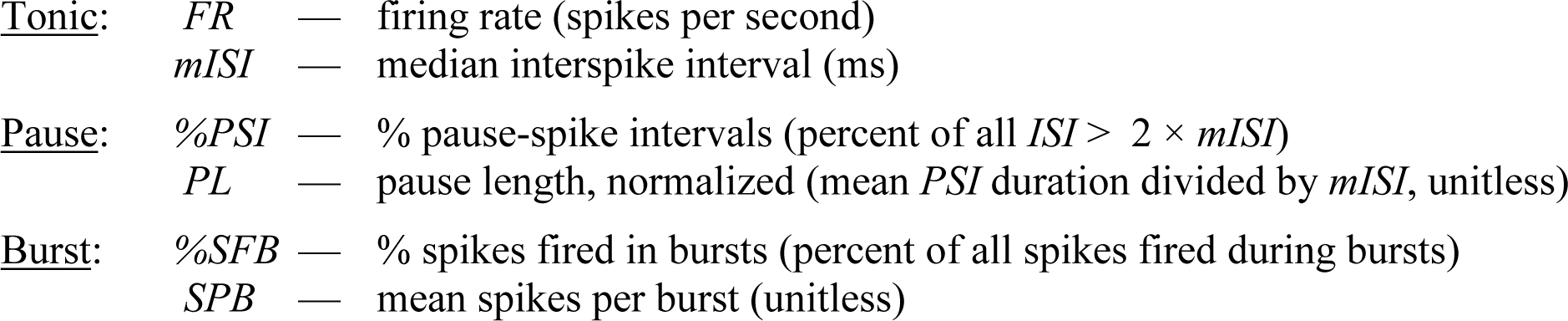

For burst metrics, a burst is defined as a sequence of 3-10 spikes in which the first *ISI* < 80 ms and subsequent *ISI* < 160 ms (*48, 49*).

Changes in a given metric, *m*, were analyzed by comparing the 15-min baseline (*m*_pre_) to a 15-min sliding window (*m*_post_) according to: *Δm*_norm_ = (*m*_post_ – *m*_pre_) / (*m*_post_ + *m*_pre_). We then plotted the time course of *Δm*_norm_ (as a function of the sliding-window time), and analyzed the steady-state *Δm*_norm_ (1-hr post-gabazine^DART^) using a two-sided permutation test (*75*). Correlations between metrics were analyzed with a Pearson’s test.**Behavior Experiments** — Adult DAT-IRES-cre mice (17 females, 20 males; 12 -16 weeks old) were anesthetized and stereotaxically injected with 400 nL of either AAV_rh10_-CAG-DIO-^+^HTP_GPI_-2A-dTomato-WPRE or AAV_rh10_-CAG-DIO- ^dd^HTP_GPI_-2A-dTomato-WPRE (2 × 10^12^ VG/mL, 100 nL per site, two tracks with two depths per track: -3.2 mm AP, ±0.5 mm ML, -5.0/-4.5 mm DV) with a custom Narishige injector. Mice were implanted with a bilateral metal cannula above the VTA (P1Tech; C235G-1.0; cut to 4 mm with a 1.0 mm spacing), which was lowered slowly to -3.75 mm. Mice were fitted with a plastic head bar adhered to the skull with OptiBond and dental cement, enabling head fixation. Mice were singly or pair housed post-surgery, in a 12-hr reverse light/dark cycle, with food and water provided *ad libitum*. Pair-housed mice were outfitted with head hats that clip to specially designed head bars (*71*) to prevent cannula damage from chewing by cage mates.

Mice were given a minimum of 9 days post-surgery for recovery and acclimation to the reverse light cycle. For the subsequent 3 days, mice were habituated to head-fixation and water restriction. Water was limited to 50-60 µL per gram of the mouse’s baseline weight per day, while dry food was provided *ad libitum*. The water restriction goal was 85% starting body weight; additional supplementary water was provided if mice dropped below 77% original body weight or did not pass a daily qualitative health assessment. Only 1 mouse was excluded for issues with water restriction health.

During behavioral sessions, mice were head-fixed (custom 3D printed clamps that fit custom head bars (*71*)) on a round plastic treadmill (Delvie’s Plastics, 8” plexiglass disk covered with silicone rubber) attached to a rotary encoder to collect rotation data (U.S. Digital H5-100- NE-S). Cue tones were played through a Z50 speaker, lick detection was collected with an infrared beam, and sucrose rewards were delivered via a Lee Company solenoid (LHDA1233315H HDI- PTD-Saline-12V-30PSI). A custom MATLAB script controlled the behavioral sessions and data collection via a National Instruments card (NI USB-6351 X Series DAQ). Behavior sessions lasted 1 hr per day for 12 consecutive days and were performed during the dark portion of the mouse’s circadian cycle. The order in which each mouse performed the task was pseudo-randomly counterbalanced.

During training sessions (days 1-10), mice were conditioned to associate cue A (2.5 kHz tone, 1.5 sec) with a 5 µL 10% sucrose-water reward. Conditioning trials were randomly interleaved with silent trials (with neither cue nor reward), enabling consistency in the trial-structure and reward-delivery quantities throughout training and testing sessions. On the final day of training (day 10) we replaced 5-6 of the silent trials with probe trials in which an unfamiliar cue B (11 kHz tone, 1.5 sec) was presented but unrewarded. Thereafter, on day 11, we infused 10 µM gabazine.7^DART.2^ + 1 µM Alexa647.1^DART.2^ dissolved in sterile aCSF; 0.6 - 0.8 µL was infused per hemisphere at a rate of 0.1 µL/min (Harvard Apparatus PhD Ultra pump using 5 µL Hamilton syringes). Following a 2 hr rest, mice resumed the original training rules for 15 min. Thereafter the rules changed: cue A was now unrewarded (extinction trials) interleaved with cue B rewarded (conditioning trials). These rules continued on day 12. Throughout the assay, mice completed 200–300 total trials daily (half cue A; half cue B or silent). Licks were allowed during the 1.5 sec tone (anticipatory licks) and the subsequent 2 sec period (retrieval licks). The inter-trial interval (ITI) was random 3 - 13 sec (from the end of the retrieval period to the start of the next cue). Licks occurring during the ITI resulted in a timeout penalty and resetting of the ITI to discourage nonspecific licking. Timeouts were never imposed for licking during a cue or retrieval period (regardless of whether the cue was rewarded or unrewarded).

Anticipatory licking (during the 1.5 sec cue) was our primary learning measure, which we quantify as the fraction of time that the infrared beam was broken during the cue. Our behavioral inclusion criteria required that mice exhibit mean cue A anticipatory licking greater than 0.2 on the 10^th^ training session (this was satisfied by 27/36 mice), and cue B probe-trial anticipatory licking less than 30% of responses to cue A (satisfied by 24/27 mice). The main-text figures include the 24 mice that met our behavioral inclusion criteria (12 ^dd^HTP, 12 ^+^HTP). The behavioral experimenter was blinded to virus condition in half of the experimental cohorts. Fig. S4c contains a total of 25 ^dd^HTP mice (13 females, 12 males) which include the same 12 ^dd^HTP mice (receiving gabazine.7^DART.2^) plus an additional 13 ^dd^HTP mice that also met behavioral inclusion criteria and had received a different infusion (blank.1^DART.2^, diazepam.1^DART.2^, or YM90K.1^DART.2^) at doses shown to have no behavioral ambient drug effects (*43*). Following the session on day 12, all mice were perfused for histological visualization of tracer^DART^ capture. No mice were excluded based on histology. All statistical comparisons were between ^dd^HTP vs ^+^HTP mice were determined using two-sided permutation tests (*75*).

### Fiber Photometry

Adult DAT-IRES-cre mice (12 females, 12 males; 12 -16 weeks old) were anesthetized and stereotaxically injected with a mixture containing pGP-AAV9-CAG-FLEX- jGCaMP8f-WPRE (5 x 10^11^ VG/mL) and either AAV_rh10_-CAG-DIO-^+^HTP_GPI_-2A-dTomato- WPRE or AAV_rh10_-CAG-DIO-^dd^HTP_GPI_-2A-dTomato-WPRE (2 × 10^12^ VG/mL) (400 nL total; 100nL per site, two tracks with two depths per track: -3.2 mm AP, ±0.5 mm ML, -5.0/-4.5 mm DV) with a custom Narishige injector. Mice were implanted with a unilateral mini metal cannula in one hemisphere above the VTA (P1Tech; C315GMN/SPC; cut to 7 mm), at a 5-10 degree angle towards the midline and lowered slowly to -3.75 mm DV. They were also implanted with an optic fiber (Doric Lenses, MFC_400/430-0.66_5mm_MF1.25_FLT) in the opposite hemisphere, at a 5-6 degree angle towards the midline and lowered slowly to -4.25 mm DV, just dorsal to the VTA. Mice were fitted with a plastic head bar adhered to the skull with OptiBond and dental cement, enabling head fixation. Mice were singly housed post-surgery, in a 12-hr reverse light/dark cycle, with food and water provided *ad libitum*.

After three weeks, mice performed the Pavlovian assay with fiber photometry recordings on every behavioral session (Tucker-Davis Technologies RZ10X; TDT Synapse software; 465 nm excitation). Mice that did not meet behavioral inclusion criteria were excluded (4 for insufficient anticipatory licking to cue A; 2 for insufficient discrimination of cue B). Prior to the first testing session on day 11, ligands were freshly dissolved in sterile aCSF to 10 µM gabazine.7^DART.2^ + 1 µM Alexa647.1^DART.2^ or 10 µM blank.1^DART.2^ + 1µM Alexa647.1^DART.2^. Given the need to achieve bilateral ligand delivery through a unilateral cannula, 0.8 nL was infused 2 or 3 times at a rate of 0.1 µL/min with 1 hour between each infusion (1.6-2.4 µL total; Harvard Apparatus PhD Ultra pump using 5 µL Hamilton syringes). Behavior proceeded 2 hr after the last infusion. Following the last session on day 12, histology was obtained to confirm jGCaMP8f expression, Alexa647^DART^ capture, and fiber placement. No mice were excluded based on histology. The behavioral experimenter was blinded to virus condition in all of the experimental cohorts.

To assess the effects of our manipulation on phasic activity, we computed:

*ΔF/F*_0_ *=* (*F – F*_0_) */ F*_0_ where

- *F* is the instantaneous fluorescence GCaMP intensity.
- *F*_0_ is the baseline fluorescence signal (1 sec interval before each cue). Consistent with photobleaching of GCaMP, a plot of the raw *F*_0_ vs trial number adhered to a double-exponential fit. We used this fit to estimate *F*_0_, thereby accounting for photobleaching while minimizing trial-to-trial noise.

We then defined time intervals as follows: (where *t* = 0 at the start of the 1.5 sec cue).

- Cue-evoked responses were time-averaged from *t* = 0.0 to 1.0 sec.
- Reward responses (full width) were averaged from *t =* 1.5 to 3.0 sec.
- Reward responses (narrow) were averaged from *t =* 1.75 to 2.25 sec.

To assess the effects of our manipulation on tonic activity, we computed: Δ(*F*_0_)_norm_ = (*F*_0,post_ - *F*_0,pre_) / (*F*_0,post_ + *F*_0,pre_) where:

- *F*_0,pre_ is averaged over days 8-10.
- *F*_0,post_ is averaged over days 11-12.

Given that all mice had met our behavioral inclusion criteria, having learned the cue A reward association, cue-evoked bursts provide a quality-control metric for GCaMP expression and fiber placement. Thus, our photometry inclusion criteria required that cue-evoked *ΔF/F*_0_ > 1, averaged over days 6-10. To confirm the appropriateness of this threshold, we performed a regression analysis against our primary metric, the early-pause *ΔF/F*_0_ (first 4 reward omissions). This regression analysis, which included all mice, confirmed a statistically significant difference between ^dd^HTP and ^+^HTP mice (*P* = 0.008, two-sided permutation slope test, **Fig. S4B**), while demonstrating the appropriateness of our inclusion threshold, below which pauses could not be reliably detected in control mice. All statistical comparisons were between ^dd^HTP vs ^+^HTP mice were determined using two-sided permutation tests (*75*).

### Histology

Mice were deeply anesthetized with isoflurane. Electrodes were briefly connected to a 9V battery (1 sec) to mark electrode positions. Thereafter, mice were fixed by transcardial perfusion of 15 mL PBS followed by 50 mL ice-cold 4% paraformaldehyde (PFA) in 0.1M PB, pH 7.4. Brains were excised from the skull, post-fixed in 50 mL of 4% PFA at 4°C overnight, then washed three times with PBS. Brains were embedded in 5% agarose and sliced along the coronal axis at 50 µm (Leica, VT1200S).

For tyrosine hydroxylase immunostaining, sections were washed in PBS before a 2 hr incubation in a blocking solution consisting of 5% goat serum, 3% bovine serum albumin, and 0.3% trition-x. Sections were then transferred to a half block solution containing 1:1000 rabbit anti-TH (PelFreez, P40101) overnight at 4°C with agitation, and then washed in 0.1M PBS containing 0.1% tween before a 4 hr incubation in a half block solution containing 1:1000 goat anti-rabbit 488 (Invitrogen, A11008). Finally, sections were washed in PBS containing tween, then PBS alone prior to mounting on glass slides.

Sections were mounted onto glass slides (VWR 48311-703) and coverslipped with Vectashield mounting medium (Vector Labs, H-1400 or H-1800). Fluorescent images (DAPI, FITC, TRITC, Cy5) were collected at 10X magnification with an Olympus VS200 slide scanner.

Cell counts were obtained using ilastik (*76*). Pixel Classification was used to predict cell versus not-cell (background tissue), then Object Classification was used on these pixel predictions to label cells as red (dTomato), green (TH+ or GCaMP), or red+green (both). Object identities were exported and used to calculate the number of cells identified in each label class across all sections from one brain. Pixel intensity analysis was performed with custom MATLAB code. For each coronal section, the VTA was manually segmented in both hemispheres. Background fluorescence was subtracted. Dye capture levels were calculated via a pixel-wise summation over 15 coronal sections. Correlations between pixel intensity and behavior were analyzed with a Pearson’s permutation test; trend lines are simple linear regressions, and shading is 95% confidence interval.

---

**Fig. S1:**
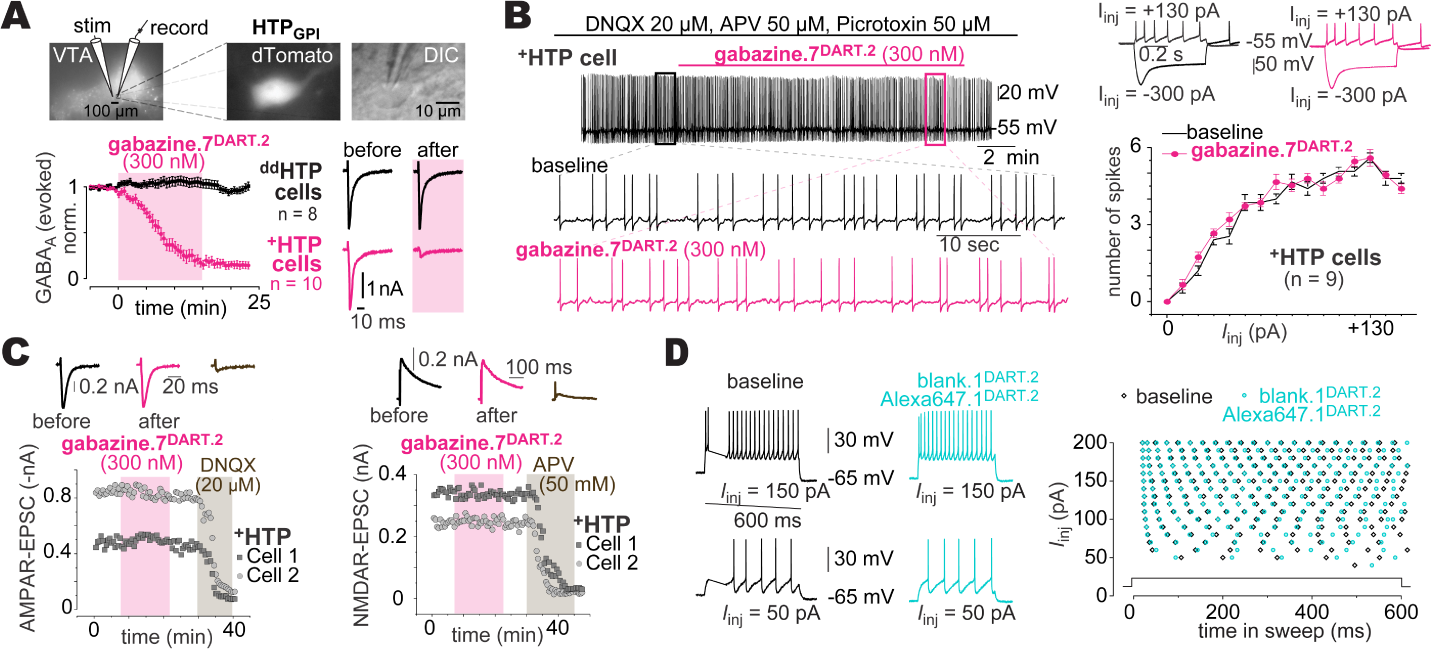
Specificity of gabazine^DART^ to the GABA_A_R A**: Gabazine.7^DART.2^ validation.** Top: evoked-IPSC configuration in VTA slice. Bottom: 300 nM gabazine.7^DART.2^ has no impact on ^dd^HTP neurons, while blocking IPSCs on ^+^HTP cells in under 15 min (86 ± 4% block). Data are mean ±SEM, cells normalized to baseline (^+^HTP: n=10 cells; ^dd^HTP: n=8 cells). Example traces to right. B**: Gabazine.7^DART.2^ and VTA_DA_ action potentials.** Left: current clamp of a VTA_DA_ neuron in the presence of picrotoxin (GABA_A_ receptor blocker), DNQX (AMPA receptor blocker), and APV (NMDA receptor blocker). Right: quantification of action potential firing as a function of injected current; performed before (black) vs after (cyan) gabazine.7^DART.2^ was tethered on each cell. Representative traces shown above. Error bars are mean ±SEM over cells (*n* = 9). C**: Gabazine.7^DART.2^ and AMPARs/NMDARs.** Left: 300 nM gabazine.7^DART.2^ has no effect on ^+^HTP neuron AMPAR-EPSCs, which are subsequently blocked by 20 µM DNQX. Example traces above. Right: 300 nm gabazine.7^DART.2^ has no effect on ^+^HTP neuron NMDAR-EPSCs, which are subsequently blocked by 50 µM APV. Example traces above. D**: Alexa647.1^DART.2^ validation.** Current clamp studies in VTA_DA +_HTP neurons with 10:1 blank.1^DART.2^ + Alexa647.1^DART.2^. No significant change was observed before vs after Alexa647.1^DART.2^ was tethered. Representative traces shown left.

**Fig. S2:**
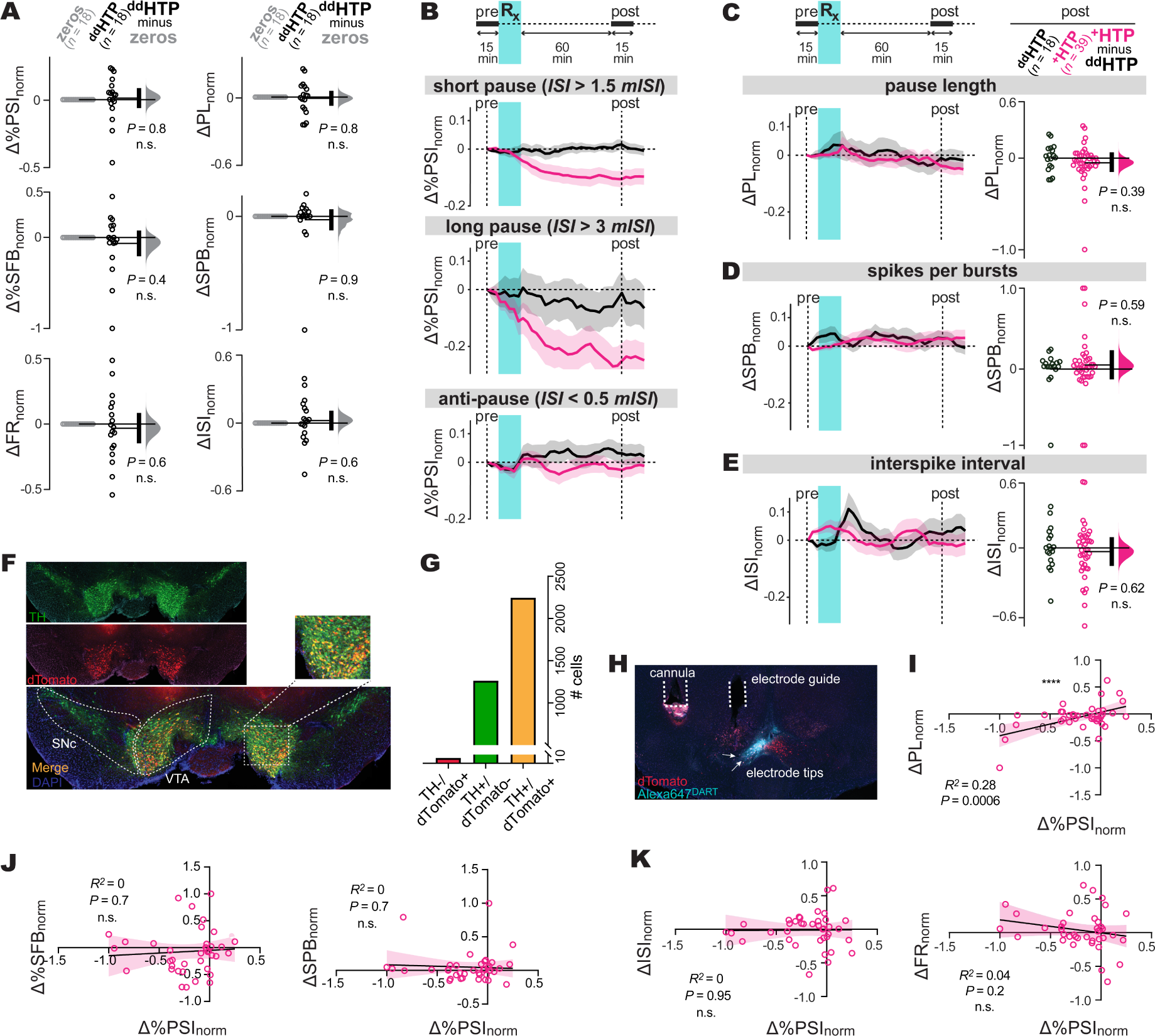
Supporting data for *in vivo* electrophysiology A**: Control ^dd^HTP metrics.** Analysis of tonic, burst, and pause stability in ^dd^HTP mice. Steady-state Δ_norm_ (1-hr post-gabazine^DART^) with individual cells (circles), group means (thin horizontal lines), mean-difference bootstrap (grey distribution), and 95% CI of the two-sided permutation test (vertical black bar), comparing ^dd^HTP cells to zero. B**: Examining robustness of pause result.** Robustness analysis of our primary pause metric, *%PSI*. The top two panels examine various “longer-than-average” metrics of pause occurrence (format as in Fig. 1d). The bottom panel examines a “shorter-than- average” metric to test for the asymmetry of effects. C-E**: Pause length, bursts, and ISI.** Analysis of *PL* (pause length), *SPB* (spikes per burst), and *mISI* (median interspike interval); format as in Fig. 1d-f. F-G**: Cell counting.** Representative histology and quantitative cell counting. Dopamine neurons (TH, tyrosine-hydroxylase, green). HTP expression (dTomato, red). Cell counting was performed from one representative brain. There were 2,244 double-labeled (dTomato^+^/TH^+^) cells. This represents 99.7% of all virus-positive cells (2,250 dTomato^+^), and 64% of all dopaminergic cells (3,507 TH^+^). H**: Electrode histology.** Post-electrophysiology histology. HTP expression (dTomato, red); ligand capture (cyan); and electrode tips (arrows). I**: Pauses vs pause length.** Correlation between *PL* (pause length) and *%PSI* from each ^+^HTP cell (circles, *n*=39), with regression ±95% CI (line and shading). Pearson’s *r*^2^ = 0.28, *P* = 0.0006. J-K**: Pauses vs other features.** Correlation between all other metrics and *%PSI*; format as above.

**Fig. S3:**
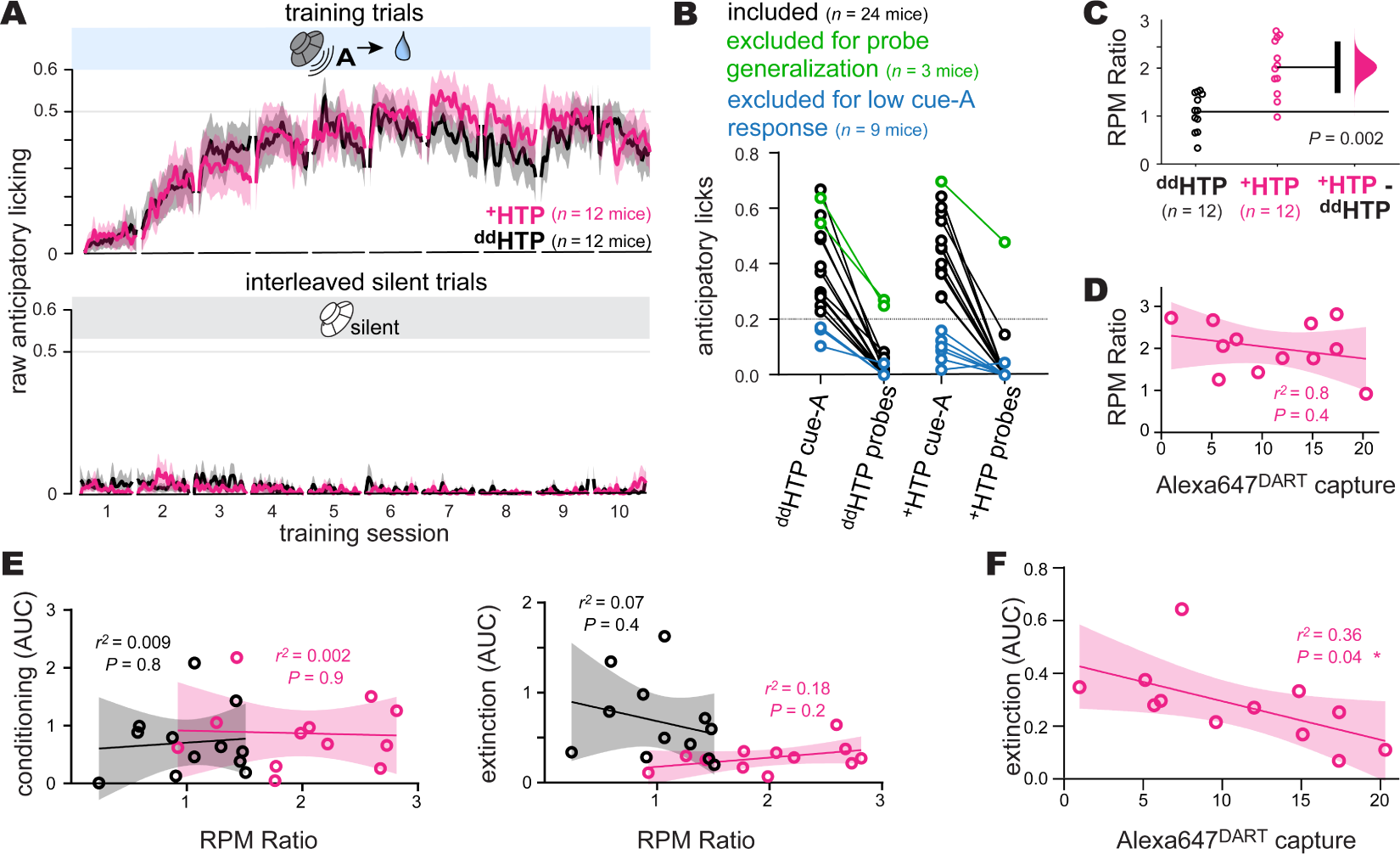
Supporting data for Pavlovian extinction and conditioning assay A**: Training sessions**. Lines and shading show anticipatory licking (fraction of time that beam is broken during the cue), mean ± SEM over mice (n = 12 ^dd^HTP; n = 12 ^+^HTP). Both ^+^HTP and ^dd^HTP mice develop robust anticipatory licking to cue A across training, while exhibiting little to no background licking during silent trials. Note that this figure shows raw (non-normalized) anticipatory licking, whereas the main-text figures display normalized anticipatory licking, with day-10 anticipatory as a constant of normalization for each animal. B**: Behavioral inclusion criteria**. We required robust anticipatory licking to cue A (raw anticipatory licking greater than 0.2) and low responsiveness to cue B probes (less than 30% of cue A anticipatory licking). A total of 9 mice (3 ^dd^HTP and 6 ^+^HTP) were excluded for lack of cue A responsiveness (blue). Of the remaining 27 mice, only 3 mice (2 ^dd^HTP and 1 ^+^HTP) were excluded for probe generalization (green). Thus 89% (24 of 27) successfully discriminated cue B. C**: Locomotion**. Ratio of the average treadmill RPM post-gabazine^DART^ (day 11-12) divided by pre-gabazineDART (day 8-10). Individual mice (circles), group means (thin horizontal lines), mean-difference bootstrap (pink distribution), and 95% CI of the two-sided permutation test (vertical black bar). As previously reported (*43*), disinhibition of VTA_DA_ neurons enhances locomotion. D**: Locomotion vs histology**. Correlation between RPM ratio and Alexa647^DART^ capture in the dorsal VTA of ^+^HTP mice (*n*=12). Mice (circles), regression ±95% CI (line and shading). Pearson’s *r*^2^ = 0.08, *P* = 0.4 indicates no significant correlation. E**: Locomotion vs reward learning**. Correlation between RPM ratio and our two measures of reward learning: conditioning AUC (left) and extinction AUC (right). Mice (circles; *n*=12 ^dd^HTP; *n*=12 ^+^HTP) and regression ±95% CI (line and shading). Pearson’s tests show no significant correlation (*r*^2^ and *P* values as indicated). F**: Extinction learning vs histology**. Correlation between extinction (AUC) and Alexa647^DART^ capture in the dorsal VTA of ^+^HTP mice (*n*=12). Mice (circles), regression ±95% CI (line and shading). Pearson’s *r*^2^ = 0.36, *P* = 0.04 indicates a significant correlation, with higher levels of target engagement corresponding to faster rates of extinction.

**Fig. S4:**
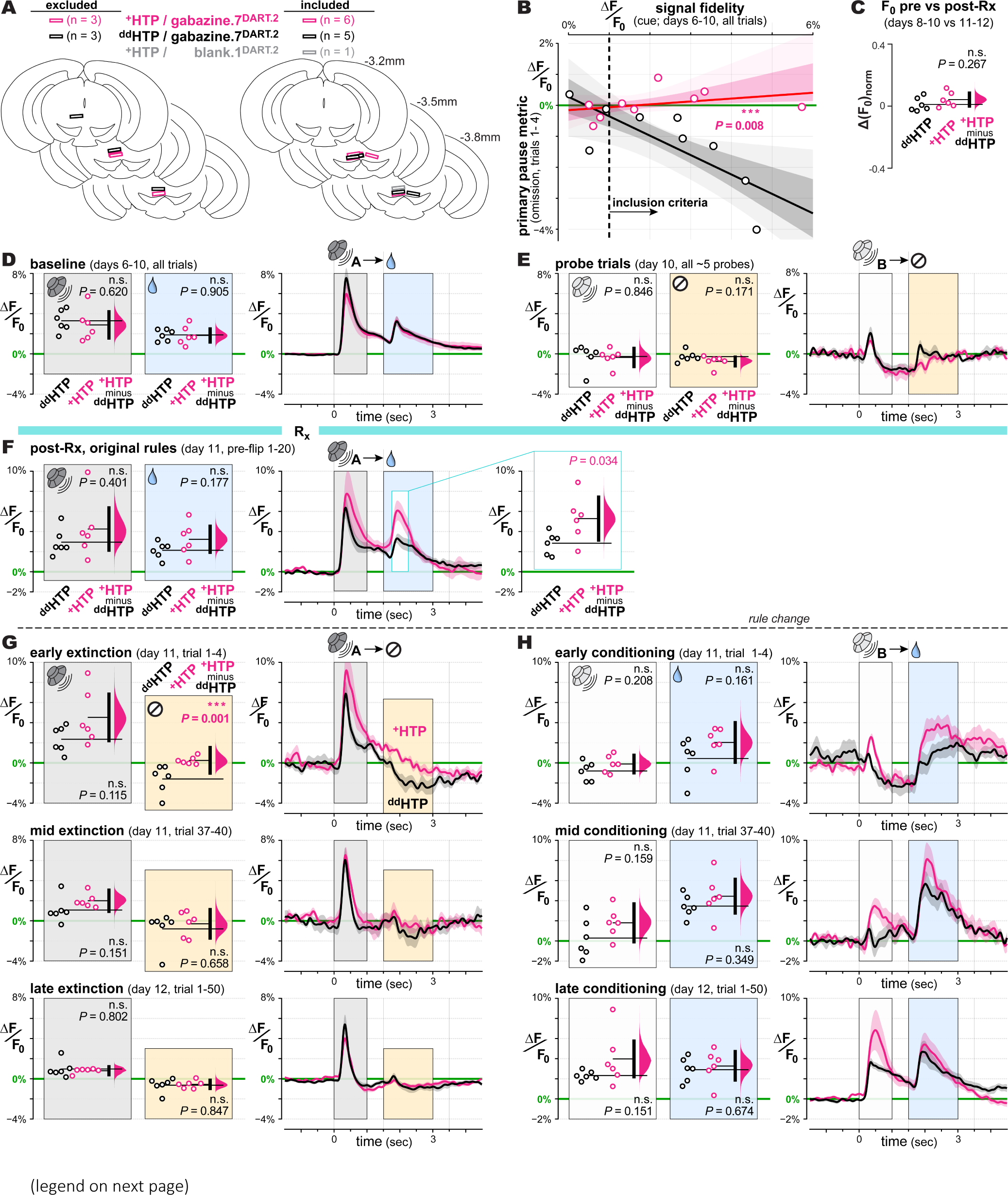
Supporting data for fiber photometry studies A**: Fiber placement**. Optic fibers were placed dorsal to the VTA and equivalently spread between control and experimental conditions. 6 mice (3 ^+^HTP/gabazine^DART^ and 3 ^dd^HTP/gabazine^DART^) were excluded based on GCaMP8f signal levels (left). 12 mice (6 ^+^HTP/gabazine^DART^, 5 ^dd^HTP/gabazine^DART^, and 1 ^+^HTP/blank^DART^) were included in further analysis (right). B**: Photometry inclusion criteria**. Signal-fidelity metric (cue-evoked burst, days 6-10) reflects GCaMP8f expression and its coupling efficiency to the fiber-optic. This pre-gabazine^DART^ signal-fidelity metric is plotted against our main post-gabazine^DART^ pause metric (omission pause, day 11 extinction trials 1-4). Data from individual mice (circles), regression fits (lines), and regression 95% and 68% CI (light and dark shading) are shown for ^dd^HTP (black, *n* = 9) and ^+^HTP (pink, *n* = 9) mice. With all data included, there is a clear statistical difference between ^dd^HTP and ^+^HTP mice (two-sided permutation slope test, *P* = 0.008). Mice above a signal- fidelity threshold (to the right of the dashed line at 1% ΔF/F_0_) are included in the subsequent analyses. C**: Tonic activity**. Changes in baseline GCaMP8f intensity on days 8-10 (F_0,pre_) vs days 11-12 (F_0,post_) are analyzed according to Δ(F_0_)_norm_ = (F_0,post_ - F_0,pre_) / (F_0,post_ + F_0,pre_). Individual mice (circles), group means (thin horizontal lines), mean-difference bootstrap (pink distribution), and 95% CI of the two-sided permutation test (vertical black bar); ^+^HTP and ^dd^HTP cells do not differ significantly (*P*=0.267). D**: Days 6-10 rewarded cue A trials.** Right: GCaMP8f responses in ^dd^HTP (black) vs ^+^HTP (pink) mice during days 6-10 (average over all cue A trials). Right panel shows the time course of ΔF/F0 mean ± SEM over mice (*n* = 6 ^dd^HTP mice; *n* = 6 ^+^HTP mice). Analysis of ΔF/F_0_ during cue (gray) and reward (yellow) is plotted in the left panel of corresponding color. Left: individual mice (circles), group means (thin horizontal lines), mean- difference bootstrap (pink distribution), and 95% CI of the two-sided permutation test (vertical black bar). ^+^HTP and ^dd^HTP mice were not statistically different during cue (0 – 1 sec interval; *P*=0.620) or reward (1.5 – 3 sec interval, *P*=0.905). Examination of an additional, narrow-time reward interval (1.75 – 2.25 sec) was also not significant (*P*=0.968; not shown). E**: Day 10 unrewarded cue B probes.** Format as in panel d; for pre-gabazine^DART^ unrewarded cue B probes. Cue-evoked signals (0 - 1 sec) were not statistically different in ^+^HTP vs ^dd^HTP mice (*P*=0.846). Signals in the unrewarded interval displayed a non-significant trend (1.5 - 3 sec; *P*=0.171), which remained non-significant over a narrow interval (1.75 – 2.25 sec; *P*=0.054; not shown). F**: Day 11 rewarded cue A trials.** Format as in panel d; post-gabazine^DART^ rewarded cue A trials (prior to rule change). Cue-evoked signals (0 - 1 sec) were not statistically different in ^+^HTP vs ^dd^HTP mice (*P*=0.401). For reward-evoked signals, ^+^HTP mice exhibited a trending increase (1.5 - 3 sec; *P*=0.117) which was weakly significant within a narrow interval (1.75 – 2.25 sec; *P*=0.034, right inset). G**: Days 11-12 cue A extinction.** Format as in panel d; post-gabazine^DART^ cue A extinction (early, middle, and late trials from top to bottom). Cue-evoked signals (0 - 1 sec) showed a non-significant trend during early (*P*=0.115) and middle (*P*=0.151) but not late (*P*=0.802) trials. Omission-pause signals (1.5 - 3 sec) were evident in ^dd^HTP mice yet absent in ^+^HTP mice during early trials (*P*=0.001). This difference was not apparent during middle (*P*=0.658) and late (*P*=0.847) trials owing to the lack of pauses in control mice. Examination of a narrow interval (1.75 – 2.25 sec) upheld these results (early *P*=0.00045; middle *P*=0.448; late *P*=0.945). H**: Days 11-12 cue B conditioning.** Format as in panel d; post-gabazine^DART^ cue B conditioning (early, middle, and late trials from top to bottom). Cue-evoked signals (0 - 1 sec) showed a non-significant trend (early *P*=0.208; middle *P*=0.159; late *P*=0.151). Reward-evoked signals displayed a similar trend for both the full (1.5 - 3 sec) interval (early *P*=0.161; middle *P*=0.349; late *P*=0.674) and narrow (1.75 – 2.25 sec) interval (early *P*=0.123; middle *P*=0.306; late *P*=0.640).

**Fig. S5:**
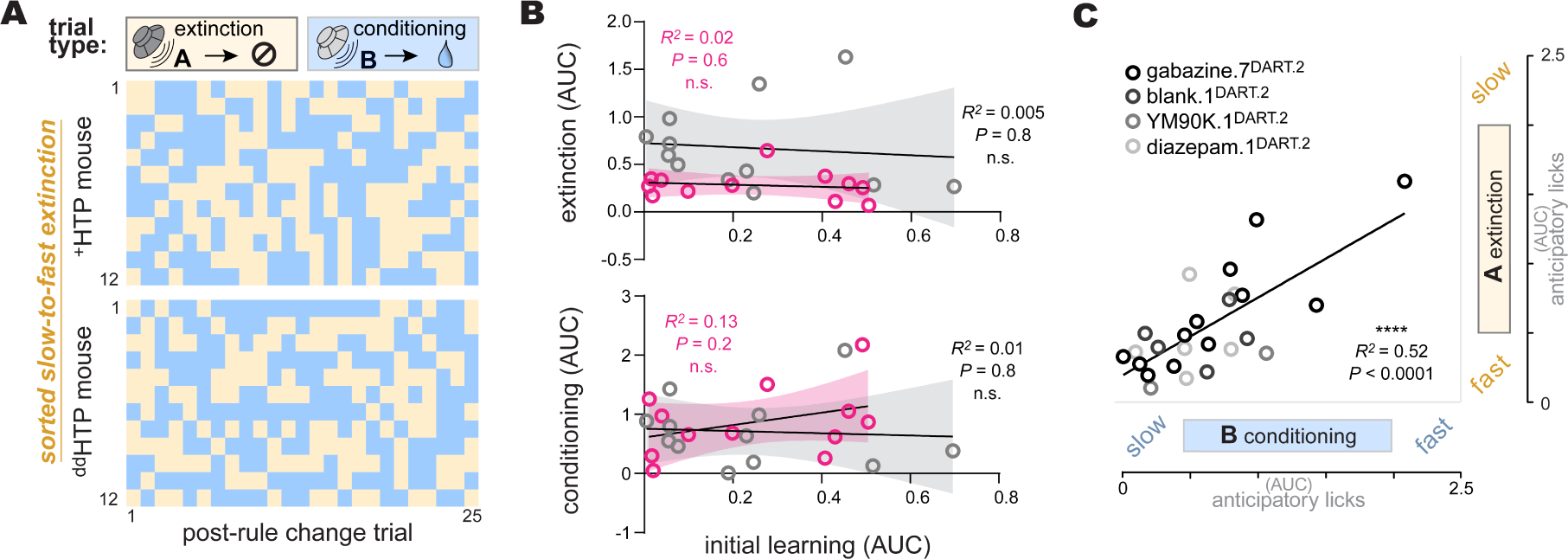
Supporting data for within-mouse behavioral correlations A**: Visual inspection of random order of interleaved trials.** Each row is one mouse, sorted by extinction from the slowest to fastest (top to bottom). Columns indicate the first 25 trials after the rule-change, with trial type indicated by color (yellow = cue A extinction, blue = cue B conditioning). No visually discernable pattern is apparent in either ^+^HTP mice (n=12) or ^dd^HTP mice (n=12). B**. Initial training vs later reward learning**. Correlation between initial learning rates (AUC over training days 1- 2) and our two measures of reward learning: extinction AUC (top) and conditioning AUC (bottom). Mice (circles; *n*=12 ^dd^HTP; *n*=12 ^+^HTP) and regression ±95% CI (line and shading). Pearson’s tests show no significant correlation (*r*^2^ and *P* values as indicated). C**: Phenotypic spectrum across pooled controls.** Conditioning-*AUC* vs extinction-*AUC* measured within- mouse. Individual mice (circles), regression fit (line), are shown for mice pooled from ongoing control experiments. All data are from ^dd^HTP mice infused with various ligands, including gabazine.7^DART.2^, blank.1^DART.2^, YM90K.1^DART.2^, or diazepam.1^DART.2^. Pearson’s *r*^2^=0
.52, *P*<0.0001. 75

**Table S1:**
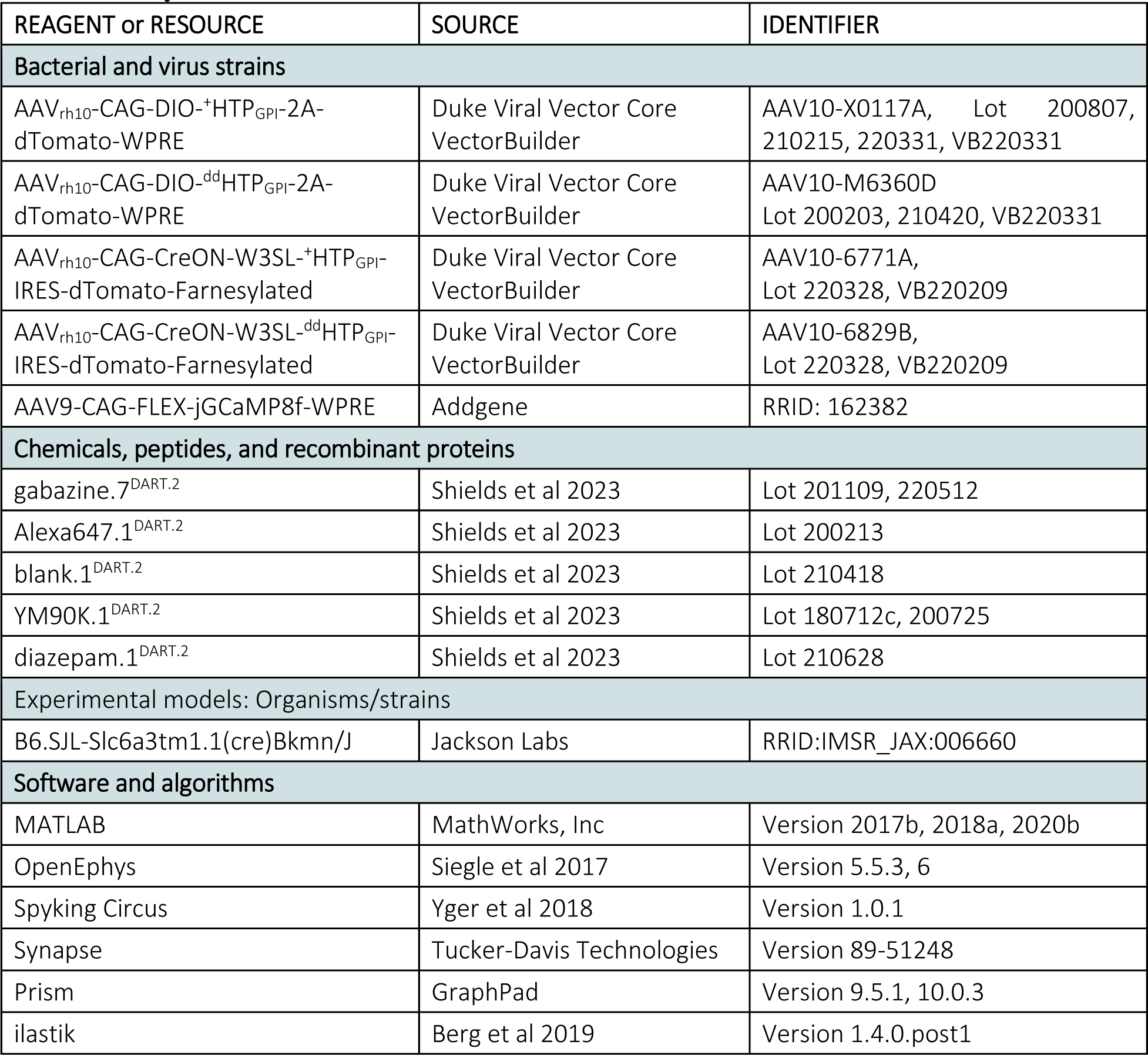
Key Resources.

| REAGENT or RESOURCE | SOURCE | IDENTIFIER |
| --- | --- | --- |
| **Bacterial and virus strains** |  |  |
| AAV<sub>rh10</sub>-CAG-DIO-<sup>+</sup>HTP<sub>GPI</sub>-2A-dTomato-WPRE | Duke Viral Vector Core VectorBuilder | AAV10-X0117A, Lot 200807, 210215, 220331, VB220331 |
| AAV<sub>rh10</sub>-CAG-DIO-<sup>dd</sup>HTP<sub>GPI</sub>-2A-dTomato-WPRE | Duke Viral Vector Core VectorBuilder | AAV10-M6360D<br>Lot 200203, 210420, VB220331 |
| AAV<sub>rh10</sub>-CAG-CreON-W3SL-<sup>+</sup>HTP<sub>GPI</sub>-IRES-dTomato-Farnesylated | Duke Viral Vector Core VectorBuilder | AAV10-6771A,<br>Lot 220328, VB220209 |
| AAV<sub>rh10</sub>-CAG-CreON-W3SL-<sup>dd</sup>HTP<sub>GPI</sub>-IRES-dTomato-Farnesylated | Duke Viral Vector Core VectorBuilder | AAV10-6829B,<br>Lot 220328, VB220209 |
| AAV9-CAG-FLEX-jGCaMP8f-WPRE | Addgene | RRID: 162382 |
| **Chemicals, peptides, and recombinant proteins** |  |  |
| gabazine.<sup>7DART.2</sup> | Shields et al 2023 | Lot 201109, 220512 |
| Alexa647.<sup>1DART.2</sup> | Shields et al 2023 | Lot 200213 |
| blank.<sup>1DART.2</sup> | Shields et al 2023 | Lot 210418 |
| YM90K.<sup>1DART.2</sup> | Shields et al 2023 | Lot 180712c, 200725 |
| diazepam.<sup>1DART.2</sup> | Shields et al 2023 | Lot 210628 |
| **Experimental models: Organisms/strains** |  |  |
| B6.SJL-Slc6a3tm1.1(cre)Bkmn/J | Jackson Labs | RRID:IMSR_JAX:006660 |
| **Software and algorithms** |  |  |
| MATLAB | MathWorks, Inc | Version 2017b, 2018a, 2020b |
| OpenEphys | Siegle et al 2017 | Version 5.5.3, 6 |
| Spyking Circus | Yger et al 2018 | Version 1.0.1 |
| Synapse | Tucker-Davis Technologies | Version 89-51248 |
| Prism | GraphPad | Version 9.5.1, 10.0.3 |
| ilastik | Berg et al 2019 | Version 1.4.0.post1 |

**Table S2.**
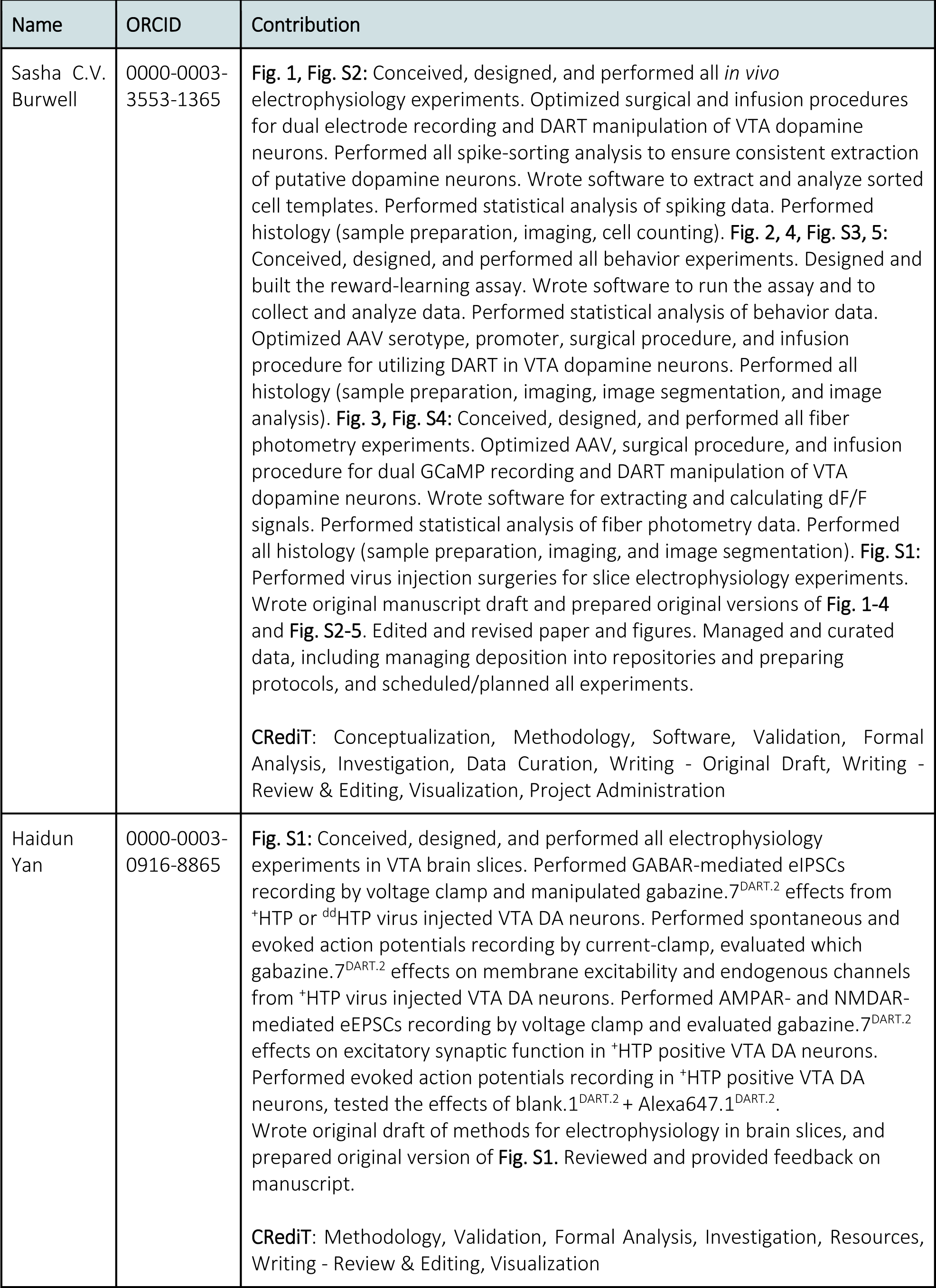
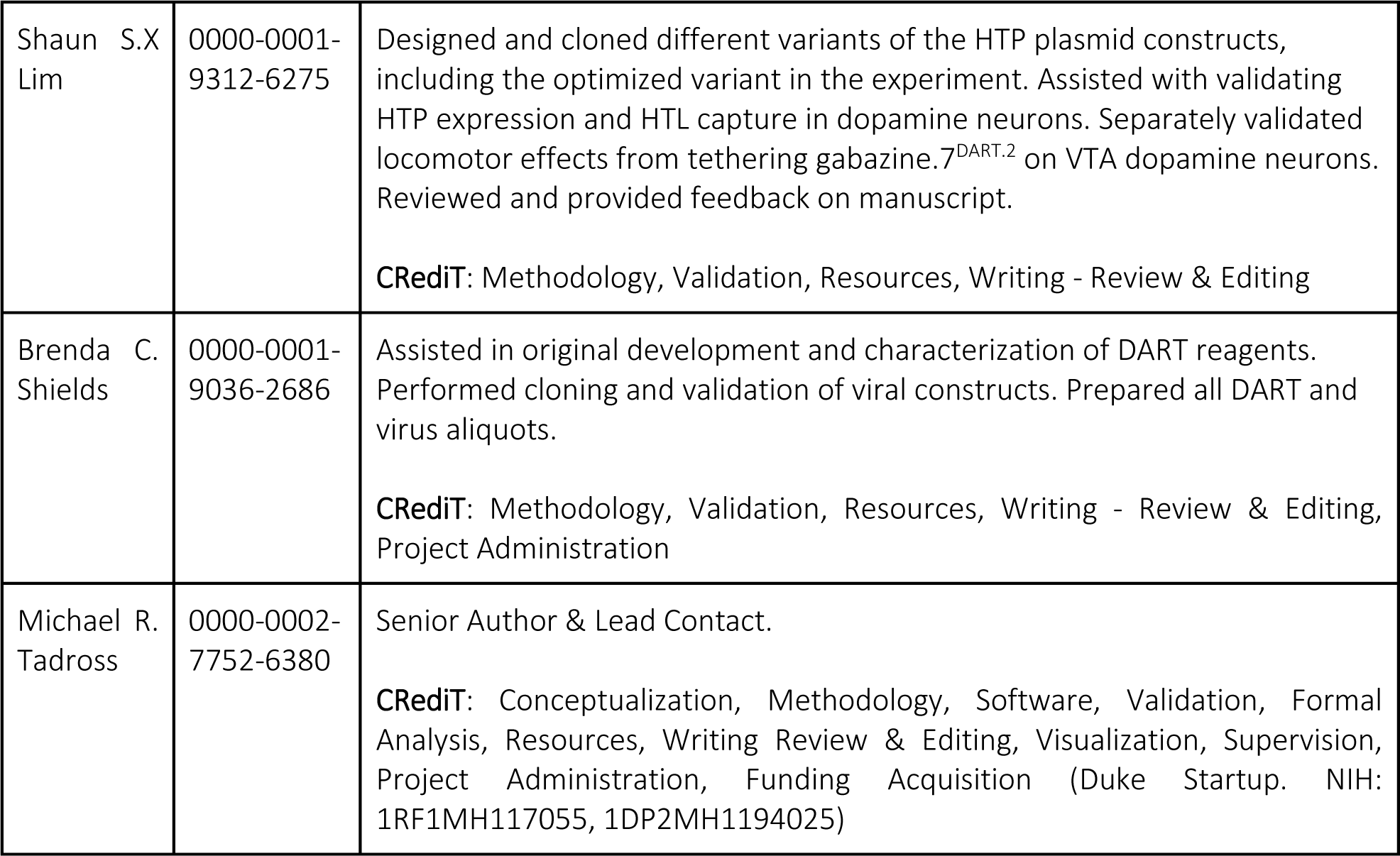
Detailed Author Contributions.

## Notes

https://doi.org/10.17504/protocols.io.j8nlk8ekdl5r/v1

https://github.com/tadrosslab/VTA_GABA_paper

https://doi.org/10.5281/zenodo.10951255

https://doi.org/10.5281/zenodo.10904059

https://doi.org/10.5281/zenodo.10903566

https://doi.org/10.5281/zenodo.10908572

https://doi.org/10.5281/zenodo.10908502

### Summary of Updates

We revised the text to improve framing and clarity. This includes an updated title and abstract. The majority of the edits were to the introduction and the discussion. All of the data, analyses, and statistics remain identical to the first version.

